# Designing DNA With Tunable Regulatory Activity Using Discrete Diffusion

**DOI:** 10.1101/2024.05.23.595630

**Authors:** Anirban Sarkar, Alejandra Duran, Yiyang Yu, Da-Wei Lin, Yijie Kang, Nirali Somia, Pablo Mantilla, Jessica Zhou, Masayuki Nagai, Ziqi Tang, Kaarina Hanington, Kenneth Chang, Peter K. Koo

## Abstract

Designing regulatory DNA with tunable and context-specific activity is a major goal in biotechnology and medicine. Deep generative models offer a promising route for sequence design, yet it remains unclear whether synthetic sequences faithfully recapitulate the motif organization and functional specificity of natural regulatory DNA. Here we present DNA Discrete Diffusion (D3), a generative model that designs regulatory DNA through an iterative nucleotide-substitution process. Across computational benchmarks, D3 improves regulatory sequence generation relative to matched diffusion baselines, producing sequences that more closely match target activity, activity distributions, and sequence composition. In K562 lentiMPRA experiments, D3-designed sequences retained measurable regulatory activity and more closely recapitulated the activity distribution of genomic regulatory sequences than matched diffusion baselines. D3 performs robustly with limited training data and generates sequences informative enough to improve predictive models when labeled data are scarce. When trained without activity labels on task-specific regulatory sequence sets, D3 learns frozen sequence representations that are predictive of enhancer activity and compare favorably to several off-the-shelf genomic language model embeddings. Analysis of the sampling process identifies reproducible phases of sequence exploration, compositional refinement, and motif-associated convergence, providing an interpretable view of how D3 constructs regulatory sequences. These results establish D3 as a practical framework for designing synthetic regulatory DNA and studying sequence features associated with context-specific regulatory activity.

## Introduction

The ability to design DNA sequences that precisely control when and where genes are expressed is a long-standing goal in biology, with broad implications for medicine, biotechnology, and gene therapy. Regulatory DNA elements such as promoters and enhancers encode these instructions, yet their function depends on short transcription factor binding motifs arranged in complex, context-dependent combinations that remain difficult to decipher^1^. High-throughput assays, including massively parallel reporter assays (MPRAs)^2,3^, STARR-seq^4^, and lentiMPRA^5^, have begun to map this regulatory logic. However, the vastness of sequence space and the strong cell-type dependence of regulatory activity continue to make prediction and design difficult.

Early approaches to regulatory sequence design relied on supervised predictive models of chromatin accessibility and gene expression^6,7^. Once trained, these models could be used to optimize sequences through gradient ascent^8,9^, simulated annealing^10^, in silico evolution^11^, motif implantation^12^, and reinforcement learning^13^, enabling several landmark demonstrations of data-driven sequence design^11–16^. However, these supervised models can exploit low-level sequence statistics, such as GC content, as predictive shortcuts^17^. Although controlling for such features is now common practice, doing so can also suppress biologically meaningful sequence properties that may contribute to regulatory function, including features related to DNA shape and flexibility^18,19^.

Deep generative models have since raised the prospect of designing synthetic regulatory DNA by learning and sampling from the distribution of functional sequences. The most widely used approaches have been genomic language models^20–22^ and, more recently, diffusion-based models^23–25^. Genomic language models are typically trained with reconstruction-based objectives, such as masked or next-token prediction^26^. These objectives may be a poor fit for aspects of metazoan regulatory DNA, where functional signal is often encoded by sparse sequence motifs embedded within otherwise weakly constrained sequence. As a result, training is dominated by background sequence reconstruction rather than the sequence features most relevant to function, a limitation that has been linked to poor performance on human regulatory prediction tasks^27,28^. Diffusion-based models offer an alternative by learning to reverse a corruption process over sequence^24^. Existing genomic diffusion models, including DDSM^23^ and DFM^25^, operate on continuous relaxations of discrete sequences. This creates a mismatch with the discrete mutational nature of regulatory DNA and makes the sampling process harder to interpret in terms of nucleotide substitutions. Discrete diffusion approaches such as D3PM^29^ have been explored in other domains, but they commonly rely on reconstruction-style objectives that may be less well aligned with sparsely encoded regulatory signal. Furthermore, evaluation remains a major challenge, as experimental testing of synthetic sequences is costly and current computational benchmarks assess sequence quality along too few dimensions to reliably distinguish models that capture regulatory grammar from those that do not.

Here, we introduce D3 (DNA Discrete Diffusion), a generative model that transforms random sequences into functional regulatory elements through an iterative mutational process operating directly in nucleotide space. D3 supports conditional generation across multiple settings, including scalar activity targets, activity profiles, and categorical cell-type labels. In addition, when trained on regulatory sequence sets without activity labels, D3 learns sequence representations that are predictive of regulatory activity. We further develop a benchmarking framework that evaluates generated sequences across functional activity, sequence composition, synthetic-real discriminability, and memorization, and we validate designed sequences experimentally using lentiMPRA in K562 cells. Across these benchmarks, D3 improves over matched continuous-relaxation diffusion baselines in most evaluation settings, and analysis of its sampling trajectory shows how sequence composition and motif-associated features emerge progressively from random sequence. Beyond sequence generation, D3 improves supervised models through data augmentation in low-data settings and supports classifier-guided generation of cell-type-conditioned enhancer sequences. Together, these results establish D3 as a useful framework for designing and evaluating synthetic regulatory DNA.

## Results

### D3: generative modeling of regulatory sequences

D3 is built on the Score Entropy Discrete Diffusion (SEDD) framework^30^, which defines diffusion directly in discrete sequence space. This allows the generative process to be represented as nucleotide substitutions rather than as diffusion over continuous relaxations of one-hot sequence encodings.

A diffusion model consists of a forward corruption process and a learned reverse process for sampling new data. In D3, these processes operate in nucleotide space {*A,C, G, T*}. Each elementary transition corresponds to a single-nucleotide substitution, but during sampling these transition probabilities are evaluated across all positions in parallel at each reverse step. The forward process follows the Jukes-Cantor substitution model^31^, which progressively randomizes each position through uniform nucleotide substitutions under a predefined noising schedule, analogous to neutral sequence evolution in the absence of selection. During training, we sample a timestep *t*, corrupt a sequence according to the forward process, and train D3 to predict a score, *s*_*θ*_ (*x, t*), for each possible nucleotide substitution (Fig. 1a). These scores are combined with the timestep-dependent noise level to compute the reverse transition probabilities. Early in sampling, high noise promotes broader exploration of sequence space, whereas later, lower-noise steps allow the learned substitution preferences to more strongly guide the sequence toward functional regulatory DNA. After training, D3 samples new sequences by iteratively applying these score-guided substitutions across all positions in parallel (Fig. 1b). Because substitutions are applied in parallel rather than autoregressively, D3 also naturally supports inpainting, in which selected positions are held fixed while the remainder are regenerated under activity constraints (see Supplementary Fig. 1).

**Figure 1.**
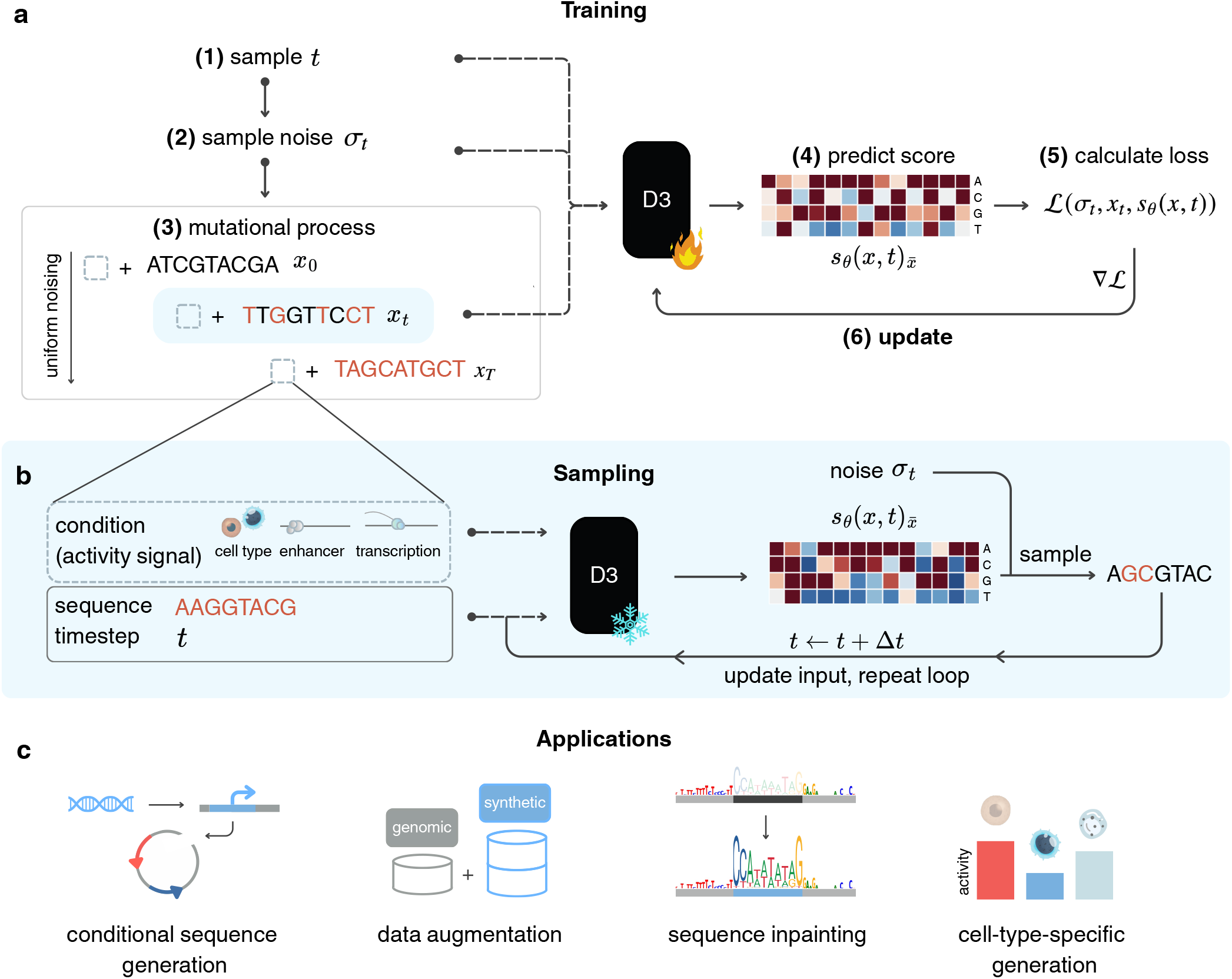
D3: a discrete diffusion framework for modeling and designing regulatory DNA. **a**, Starting from real regulatory sequences *x*_0_ and a timestep *t*, D3 applies a forward noising process that progressively corrupts DNA via uniform nucleotide substitutions. The model learns a score *s*_*θ*_ (*x, t*) that estimates density ratios between adjacent noised sequences *x*_*t*_ and *x*_*t*−1_ by minimizing the diffusion-weighted denoising score entropy loss. For conditional training, regulatory activities (multivariate profiles, scalars, or categorical variables) are concatenated with the input sequences. **b**, D3 uses learned reverse transitions to iteratively denoise random sequences *x*_*T*_ into regulatory-like sequences by predicting nucleotide substitutions at all positions and timesteps. **c**, D3 supports analyses of regulatory sequence–function relationships and applications such as data augmentation (generating plausible regulatory sequences to enrich sparse training data), sequence inpainting (filling local sequence contexts while preserving fixed sequence positions), and cell-type-conditioned generation.

SEDD is well suited for regulatory sequence modeling because it combines stable training, tractable likelihood-based learning, and flexible conditional generation within a single framework. In D3, the framework supports multiple forms of conditional generation, including scalar activity targets, activity profiles, and categorical cell-type labels. We implemented D3 with both convolutional (D3-Conv) and transformer (D3-Tran) backbones and evaluated both throughout (see Methods).

### D3 improves regulatory sequence generation across benchmarks

We evaluated D3 on three conditional generation tasks: designing human promoter sequences to match a target CAGE-seq transcription initiation profile^23^, designing Drosophila enhancers to match STARR-seq activity^32^, and designing human cis-regulatory sequences to match lentiMPRA activity measured in K562 and HepG2 cells^33^. We assessed generation quality computationally across five complementary dimensions: conditional generation fidelity, label shift, covariate shift, discriminability, and memorization (Fig. 2a–e; Supplementary Table 1). Because experimental measurements are available only for a subset of designed sequences, most generation metrics use independently trained oracle models as functional readouts. These metrics test consistency with learned sequence-function models rather than biological activity directly and are therefore interpreted together with the K562 lentiMPRA validation. Comparisons were made against DFM^25^ using the original convolutional configuration as the representative DFM baseline, together with a strictly matched transformer architecture as an architectural control across all benchmarks.

**Figure 2.**
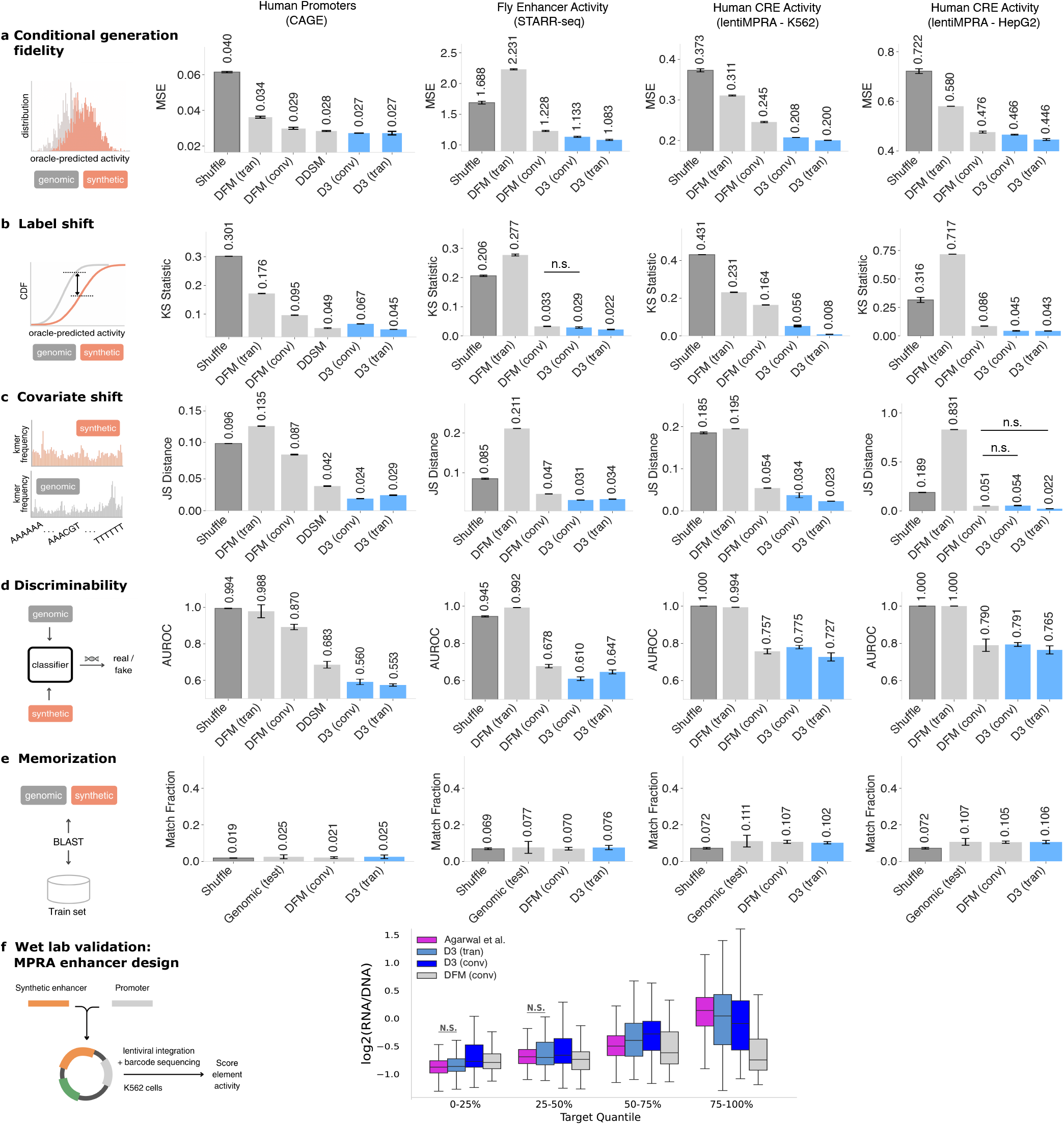
Evaluation of sequence generation quality. **a–d**, Distributional comparisons between genomic and synthetic sequences across human promoters measured by CAGE-seq (*n* = 7,497 test sequences), fly enhancers measured by STARR-seq (*n* = 41,186), and human cis-regulatory sequences measured by lentiMPRA in K562 (*n* = 39,340) and HepG2 (*n* = 24,596). Error bars represent one standard deviation across five independent sampling experiments from a single trained model. **a**, Conditional generation fidelity, measured as the MSE between oracle-predicted activity of synthetic sequences and matched target activity values. Oracle models were Sei, DeepSTARR, and MPRALegNet for CAGE-seq, STARR-seq, and lentiMPRA, respectively. **b**, Label shift, measured as the Kolmogorov–Smirnov (KS) statistic between oracle-predicted activity distributions of synthetic and genomic sequences. **c**, Covariate shift, measured as the Jensen–Shannon distance between 6-mer spectra. **d**, Discriminability, measured as the AUROC of a classifier trained to distinguish synthetic from genomic sequences. **e**, Longest contiguous exact match fraction between synthetic or reference sequences and the training set, computed with BLASTN. Dinucleotide-shuffled and held-out genomic sequences serve as reference baselines. **f**, Experimentally measured K562 lentiMPRA activity (log_2_(RNA*/*DNA)) for genomic and synthetic cis-regulatory sequences grouped by oracle-predicted activity quantile. Genomic sequences from Agarwal et al. were grouped into 0–25%, 25–50%, 50–75%, and 75–100% quantile bins (*n* = 333,326,342,325); synthetic sequences were generated with matched oracle-predicted activity targets and assigned to the same bins: D3-Tran (*n* = 54,54,52,120), D3-Conv (*n* = 54,54,54,120), and DFM-Conv (*n* = 50,56,60,120). n.s. denotes pairwise Mann–Whitney U tests versus Agarwal et al. within each bin with *p* ≥ 0.05.

Across datasets, D3-Tran provided the most consistent improvements across the evaluated metrics, while D3-Conv also improved over matched DFM baselines in most settings. The gains were most consistent for conditional generation fidelity and label shift, with additional improvements in sequence composition and synthetic-real discriminability depending on dataset and architecture. Conditional generation fidelity, measured as the mean squared error (MSE) between the oracle-predicted activity of each generated sequence and that of its matched real target sequence, was lowest for D3-Tran and D3-Conv on all benchmarks (Fig. 2a), indicating more accurate recovery of the target regulatory activity. Label shift, quantified by the Kolmogorov–Smirnov statistic comparing oracle-predicted activity distributions of generated and real sequences, was lower for D3-Tran than for matched DFM baselines across all datasets, indicating better recovery of the activity distribution (Fig. 2b). Covariate shift, quantified by Jensen–Shannon distance between 6-mer spectra of generated and genomic sequences, was lowest for D3-Tran across the lentiMPRA benchmarks and lowest for D3-Conv on the fly enhancer and human promoter benchmarks, indicating improved agreement with genomic sequence composition in most settings (Fig. 2c). Discriminability, assessed using a classifier trained to distinguish synthetic from natural sequences, was closer to random guessing for D3-generated sequences than for DFM in several settings, indicating that D3 samples were generally harder to separate from genomic regulatory sequences (Fig. 2d). The K562 lentiMPRA benchmark was a partial exception, where D3-Conv showed higher discriminability than DFM-Conv; D3-Tran outperformed both. Finally, a memorization test based on BLASTN homology search showed that the longest contiguous exact match fraction between generated and training sequences remained below 0.15 for the evaluated synthetic sequence sets (Fig. 2e), arguing against simple reproduction of training sequences.

We measured select genomic cis-regulatory sequences from Agarwal et al.^33^ spanning oracle-predicted activity quantiles together with synthetic sequences generated to match target activities from the same quantile bins. This experiment tested whether synthetic designs assigned to oracle-defined activity bins produced measured activities comparable to genomic regulatory sequences from the same bins. Among the diffusion models evaluated, D3-Tran most closely recapitulated the measured activity distribution of genomic sequences within the oracle-defined quantile bins (Fig. 2f).

### D3 representations are predictive of cis-regulatory activity

Recent benchmarks have shown that genomic language models, despite their scale, can underperform even simple one-hot encoding baselines on regression-based tasks of human regulatory activity when used without fine-tuning^27,34^. These results highlight persistent limitations in learning functional cis-regulatory signal from sequence alone. We therefore asked whether D3, trained on regulatory sequences without activity labels, could provide informative frozen sequence representations for downstream activity prediction.

We evaluated this using the lentiMPRA benchmark of Tang et al.^27^, which tests frozen sequence embeddings for predicting regulatory sequence activity in K562 and HepG2 cells measured by lentiMPRA. A fixed downstream CNN served as the probe, taking either frozen pretrained sequence representations or one-hot sequence input to predict activity (Fig. 3a). The original benchmark includes embeddings from Nucleotide Transformer^21^, DNABERT2^20^, HyenaDNA^35^, and GPN^36^, along with supervised baselines including SEI^37^, Enformer^38^, a one-hot CNN, and a ResNet trained on one-hot sequences^39,40^. We further expanded the benchmark to include Evo2^22^ in 7B and 40B versions, as well as D3 embeddings.

**Figure 3.**
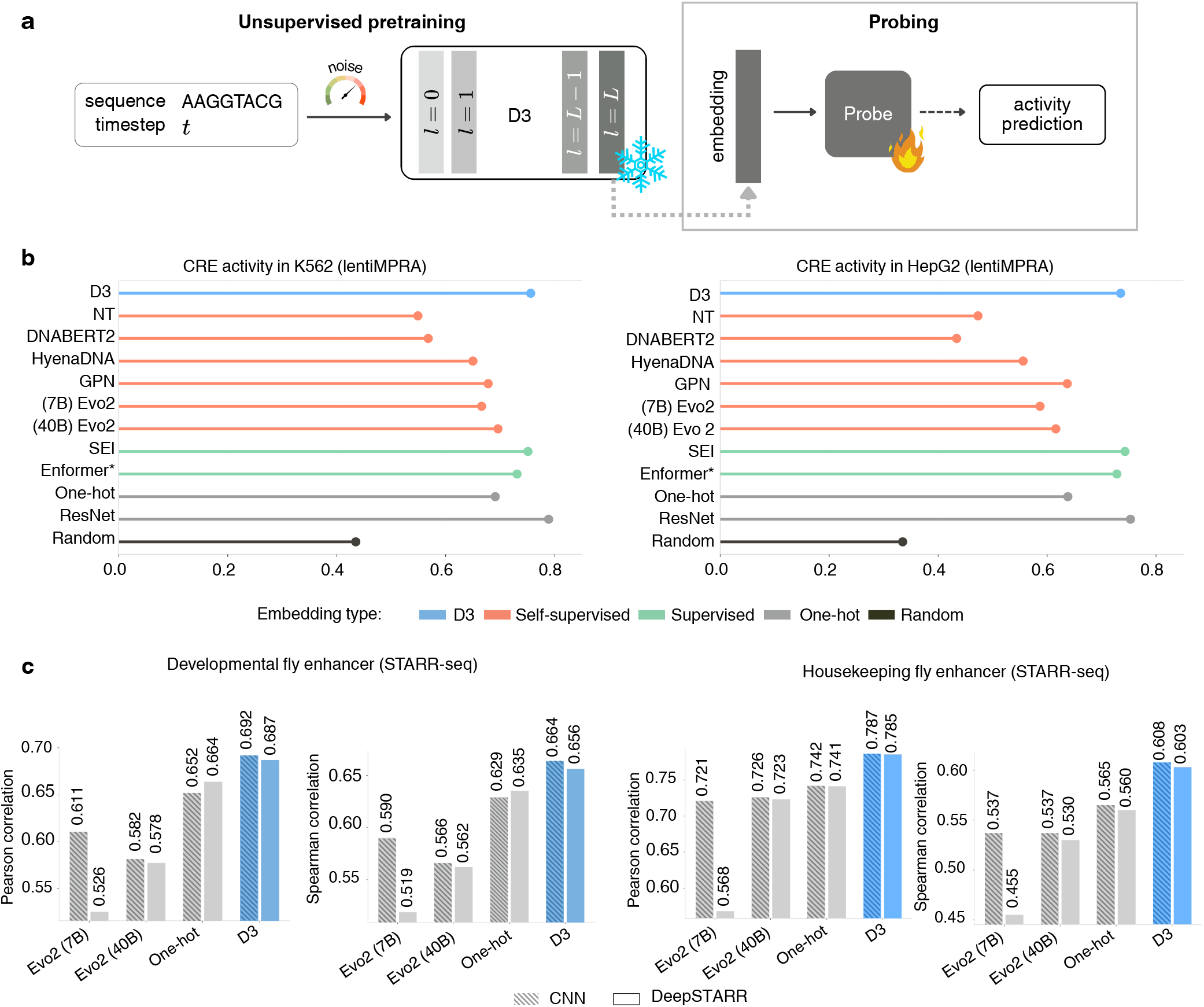
D3 learns representations predictive of regulatory sequences. **a**, Schematic of the frozen-embedding probing setup. Frozen embeddings are extracted from the final layer of the D3 score network evaluated at low noise, and passed as input to a fixed downstream CNN trained to predict regulatory activity. **b**, Pearson correlation between predicted and measured lentiMPRA activity in K562 and HepG2 cells, for frozen embeddings from D3 and a panel of genomic language models (Nucleotide Transformer, DNABERT2, HyenaDNA, GPN, Evo2-7B, Evo2-40B) and supervised baselines (SEI, Enformer, one-hot CNN, ResNet). **c**, Pearson correlation between predicted and measured STARR-seq enhancer activity in developmental and housekeeping contexts in *Drosophila* S2 cells, for D3, Evo2, and one-hot sequence representations used as input to linear, MLP, CNN, and DeepSTARR probing architectures.

In this frozen-feature benchmark, task-specific D3 embeddings outperformed the tested off-the-shelf genomic language model embeddings, including transformer-based (Nucleotide Transformer, DNABERT2), state-space (HyenaDNA, Evo2), and convolutional (GPN) architectures, in both cell types (Fig. 3b). D3 also outperformed SEI and Enformer embeddings in K562 and trailed them on HepG2 by a small margin. Because D3 was trained without activity labels, these results indicate that score-based diffusion pretraining on task-specific regulatory sequence sets can learn functionally informative representations without activity-supervised fine-tuning. Well-tuned supervised models such as ResNet still set the upper bound, but D3 substantially narrowed the gap using a simple probing model.

Next, we compared D3, Evo2, and one-hot sequence representations on fly enhancer activity prediction using linear, MLP, CNN, and DeepSTARR probing models across developmental and housekeeping tasks (Fig. 3c). D3 embeddings yielded higher Pearson and Spearman correlations than Evo2 embeddings and one-hot inputs across the displayed tasks and probe architectures. This result is notable because Evo2 is a large foundation model trained on genomic sequences across the tree of life, including the fly genome, yet still did not improve over the one-hot baseline in this setting. Both models were used as frozen feature extractors without regulatory activity labels, differing mainly in pretraining scope: D3 on the sequences from this dataset, Evo2 on genomes across the tree of life. Under this frozen-feature setup, the smaller, dataset-scoped score-based diffusion model produced more predictive representations for these enhancer activity benchmarks than the much larger broadly pretrained foundation model (92M versus 40B parameters).

### D3-designed sequences are informative for data augmentation in low-data regimes

The number of naturally occurring regulatory sequences in any given cellular context is fundamentally limited by genome size. Unlike in many machine learning domains, this imposes a hard ceiling on the amount of natural genomic data available for training supervised predictors of regulatory activity. Generative models offer a potential way to partially mitigate this constraint by producing synthetic regulatory sequences that can be used for data augmentation (Fig. 4a). We tested whether D3 could improve supervised prediction in this way using the fly enhancer STARR-seq dataset^32^.

**Figure 4.**
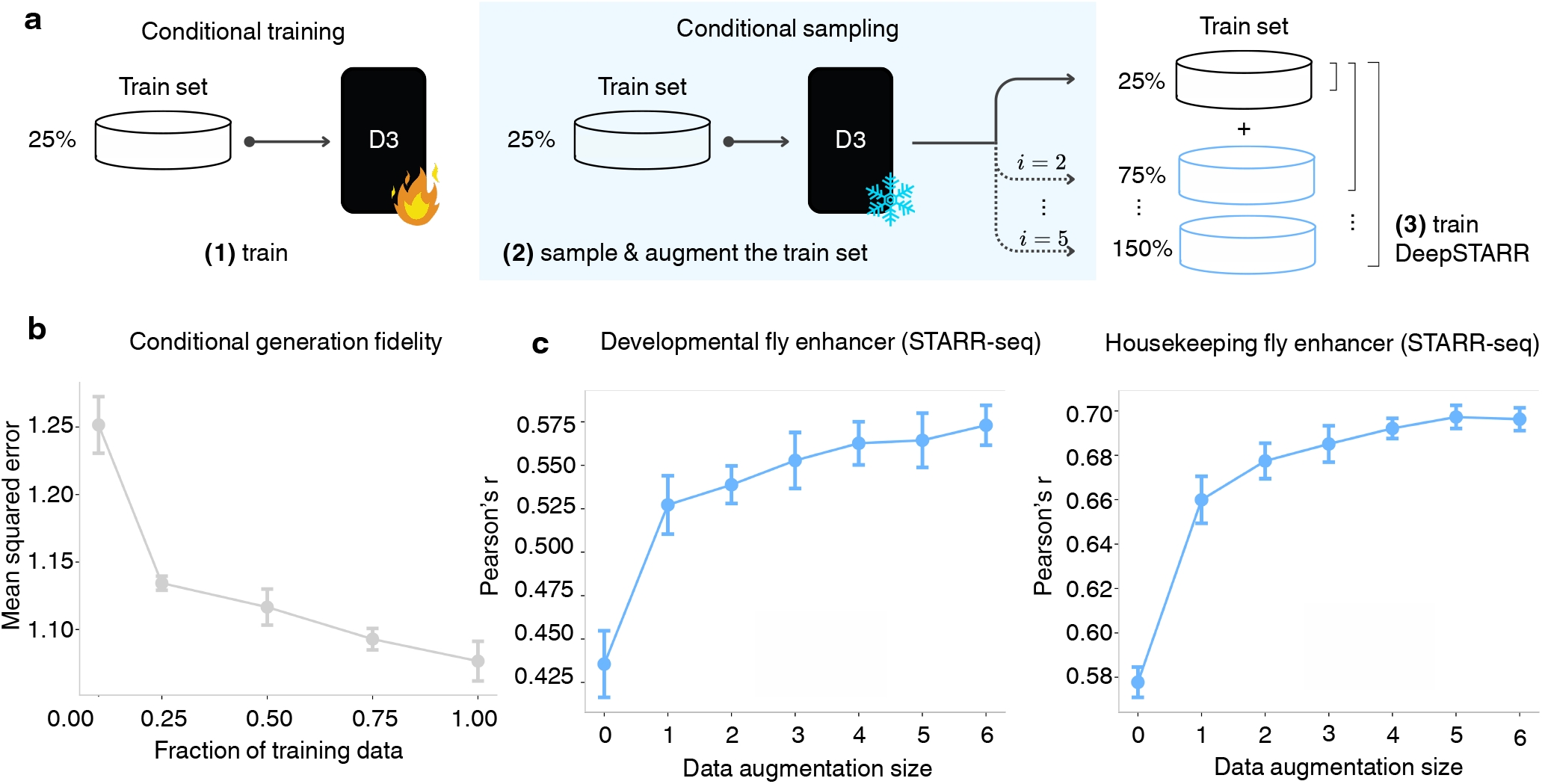
D3-generated sequences improve predictive model performance. **a**, Schematic of the data augmentation strategy using 25% of the STARR-seq training set as an example. A D3 model is trained on the subset and used to generate synthetic sequences conditioned on the subset activity values. This process augments the data size by a multiple of the subset. A DeepSTARR model is then trained on the augmented dataset and evaluated on the held-out test set. **b**, Conditional generation fidelity for D3 models trained on 10%, 25%, 50%, and 75% subsets of the STARR-seq training set. Generated sequences were conditioned based on matched test-set activity values. Error bars represent one standard deviation across five sampling experiments. **c**, Plot of performance for DeepSTARR models trained on the 25% subset augmented with D3-generated sequences at increasing augmentation sizes (1×–6× the real subset size) for developmental and housekeeping tasks. Pearson’s *r* between predicted and measured enhancer activity on the STARR-seq test set. Error bars represent one standard deviation across three independent training runs.

A prerequisite for data augmentation is that D3 itself remain effective in low-data settings. We therefore trained separate D3 models on 10%, 25%, 50%, and 75% subsets of the full training set and evaluated generation fidelity by conditioning on activity values from held-out test sequences. Generation fidelity improved substantially from 10% to 25% of the training data and continued to improve more gradually as additional data were included (Fig. 4b). This suggests that, in this dataset, D3 requires more than the smallest training subset to learn sequence features sufficient for reliable conditional generation. Above that threshold, D3-generated sequences retained sufficient conditional fidelity to test whether they could support downstream data augmentation.

We then trained D3 on the 25% subset and generated synthetic sequences conditioned on the subset’s activity values. These sequences were used to augment DeepSTARR training at increasing synthetic-to-real ratios (1x–6x, corresponding to up to six synthetic sequences per real sequence) (Fig. 4a). Augmentation consistently improved Pearson correlation on both developmental and housekeeping tasks, with gains saturating by a 5x ratio and no evidence of performance decline as additional synthetic data were added (Fig. 4c). A similar trend was also observed with a 10% subset, but with lower overall performance (Supplementary Fig. 2). Because D3 was trained on the same limited subset used for augmentation, these gains suggest that the generated sequences provided useful additional training examples for the downstream predictor. Together, these results show that D3-generated sequences can improve downstream supervised prediction in this low-data STARR-seq setting, supporting further evaluation of D3 as a source of synthetic training data when labeled regulatory sequences are scarce.

### D3 enables classifier-guided cell-type-conditioned enhancer generation

A central goal in synthetic biology and gene therapy is the design of enhancer sequences with activity patterns restricted to specific cellular contexts. As a computational test of this capability, we evaluated whether D3 could generate enhancer sequences associated with specified brain cell-type labels inferred from single-nucleus ATAC-seq, using a pretrained enhancer classifier as the readout. Whereas the conditional generation experiments above used scalar or profile-valued activity targets, this benchmark considers categorical conditioning, in which the model is asked to generate sequences for a specified cell-type label rather than a continuous activity value.

We adopted the fly brain enhancer benchmark introduced by Stark et al.^25^, based on the dataset of Janssens et al.^41^, which comprises 104,000 500-bp enhancer sequences annotated with binary activity labels across 81 brain cell types inferred from single-nucleus ATAC-seq^14^. To condition on categorical cell-type labels, we used classifier-free guidance (CFG), in which D3 was trained with conditioning labels randomly dropped from a subset of examples so that it learned both conditional and unconditional scores. At sampling time, these scores were combined to steer generation toward a specified cell type, enabling a single trained model to support all 81 cell types without retraining (Fig. 5a).

**Figure 5.**
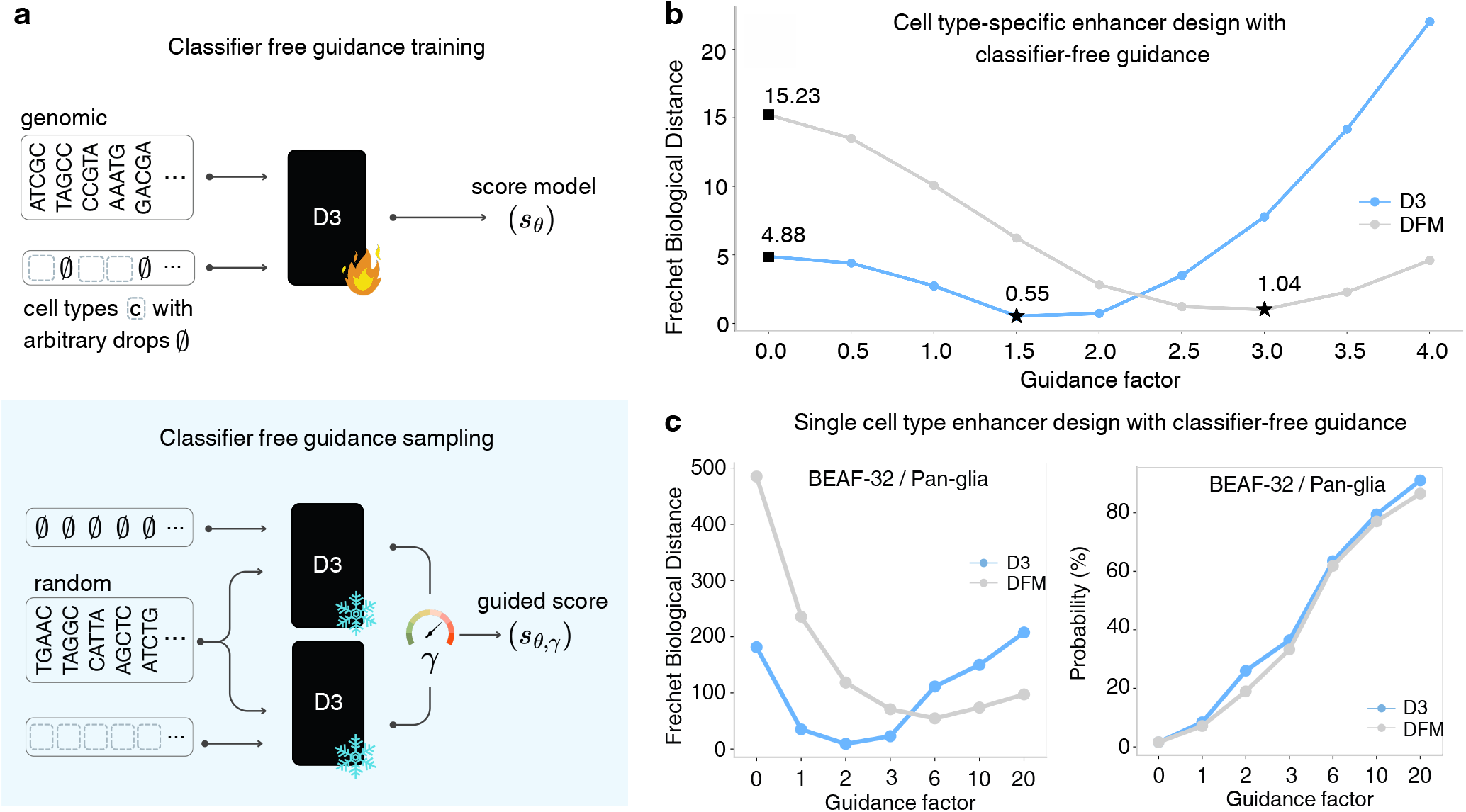
D3 generates cell-type-conditioned enhancer sequences through classifier-free guidance. **a**, Schematic of classifier-free guidance training and sampling. D3 is trained with random condition dropout, so the same sequences are presented either with their cell-type labels or with the conditioning signal removed, allowing the model to learn both conditional and unconditional score estimates. At sampling time, the two estimates are interpolated at a specified guidance strength to steer generation toward a target cell type. **b**, Fréchet Biological Distance (FBD) between generated and held-out real sequences as a function of CFG guidance strength, for D3 and DFM. Sequences are generated conditioned on 81 brain cell-type labels from the *Drosophila* brain enhancer dataset (104,000 500-bp sequences with binary activity labels across 81 cell types inferred from single-nucleus ATAC-seq). FBD is computed in the embedding space of a pretrained *Drosophila* brain enhancer classifier; lower values indicate closer alignment to the real sequence distribution. **c**, FBD (left) and target cell-type classification probability (right) assigned by a pretrained enhancer classifier to sequences generated for cell type 2 (BEAF-32 / Pan-glia) as a function of guidance strength for D3 and DFM.

We evaluated generation quality using the Fréchet Biological Distance (FBD) metric introduced by Stark et al.^25^, which measures similarity between generated and real sequences in the embedding space of a pretrained enhancer classifier, with lower values indicating closer alignment to the natural sequence distribution. D3 achieved lower FBD than DFM in the unguided setting and reached a lower minimum FBD under moderate guidance (Fig. 5b). Without guidance, D3 achieved an FBD of 4.88 versus 15.23 for DFM. With guidance, D3 reached an optimum of 0.55 at a guidance strength of 1.5, compared to DFM’s best of 1.0 at a guidance strength of 3. To assess whether generated sequences contained classifier-recognized cell-type-associated features, we measured how strongly a pretrained enhancer classifier assigned generated sequences to their intended cell type across three representative labels. At moderate guidance strengths, D3 achieved lower FBD while maintaining comparable or higher target-cell classification probabilities relative to DFM (Fig. 5c for cell type BEAF-32 / Pan-glia). FBD-vs-guidance curves and target-class probabilities for two additional cell types are shown in Supplementary Fig. 3. These results indicate that D3 better balances target-label steering with preservation of the learned enhancer sequence distribution.

As an additional qualitative demonstration of D3’s flexibility, we used inpainting to regenerate sequence context around fixed motif anchors. In this setting, selected positions are held fixed while the remaining positions are resampled under an activity constraint. For fly enhancer sequences, high-activity conditioning generated sequence context with attribution patterns resembling known activating motifs (DRE, Twist, Dorsal), whereas low-activity conditioning produced attribution patterns consistent with ttk repressor signatures and AAAGA 6-mer patterns (Supplementary Fig. 1). These examples illustrate how D3 can alter surrounding sequence context while preserving specified sequence anchors, without retraining the model.

### Motif-associated sequence features emerge progressively during D3 sampling

To characterize how D3 constructs regulatory sequences during sampling, we analyzed 1,000 trajectories for sequences conditioned on K562 lentiMPRA activity across 128 reverse-diffusion steps. Although sampling proceeds under a single continuous schedule, oracle-predicted activity follows a reproducible sigmoid-like trajectory across timesteps: sequences conditioned on high, medium, and low activity are initially indistinguishable, diverge around *t* ≈ 31, and plateau by *t* ≈ 71 (Fig. 6a, top). A complementary measure of compositional convergence, based on Jensen–Shannon distance between the *k*-mer spectra of partially sampled and genomic sequences, traces a similar decaying sigmoid (Fig. 6a, middle): it is highest early in sampling, when sequences are still undergoing broad compositional reorganization, and then decreases over time as the generated sequences become more structured and approach the data distribution. We used these transitions to define three empirical sampling stages for analysis: exploration (*t* = 0–31), consolidation (*t* = 31–71), and stabilization (*t* = 71–127).

**Figure 6.**
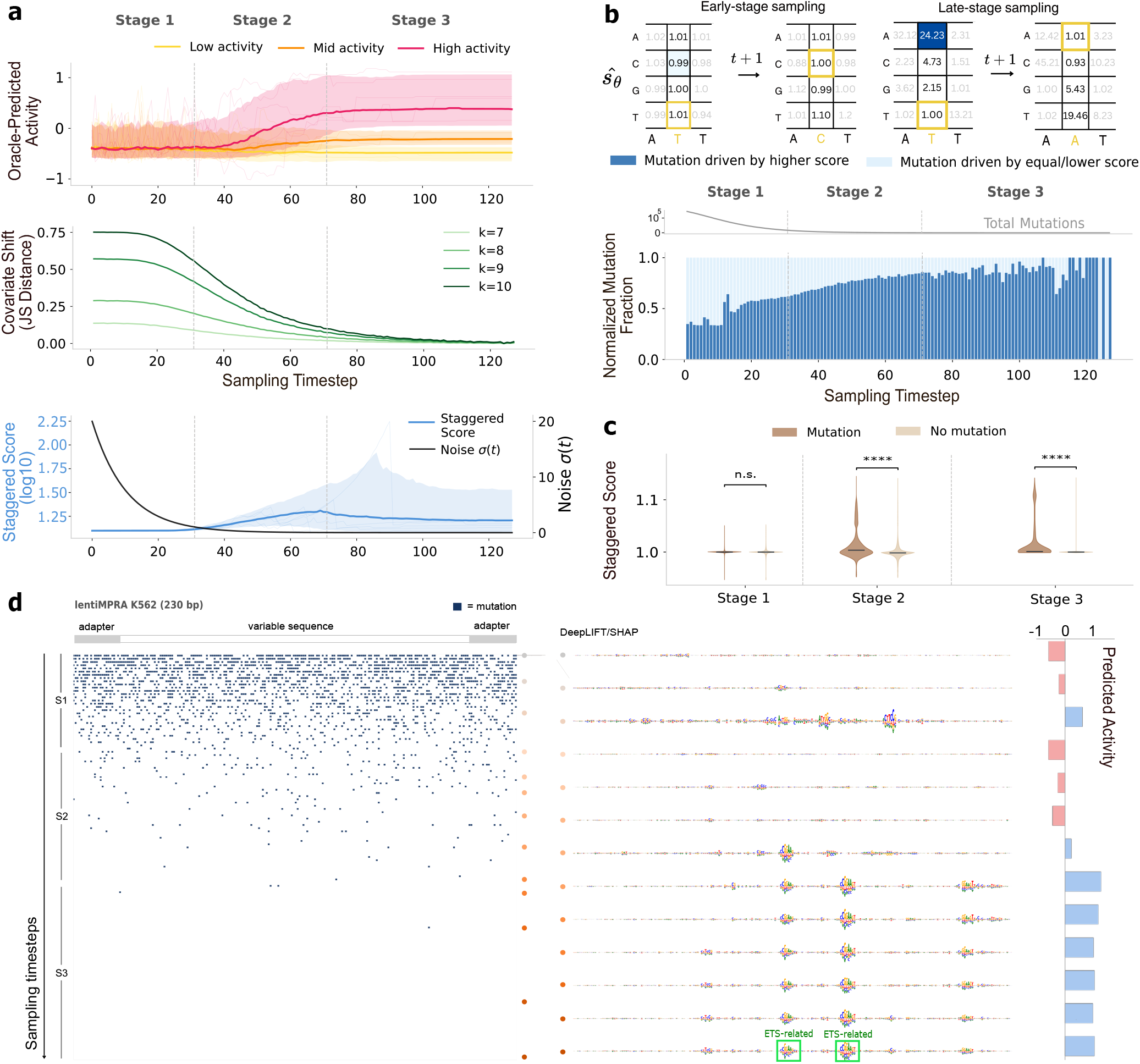
Dynamics of the discrete diffusion sampling process. **a**, Top: oracle-predicted regulatory activity (mean ± standard deviation, *n* = 1000) across 128 sampling timesteps, averaged over sequences within each activity conditioning bin (high, medium, low); thin lines show individual sample trajectories. Middle: compositional convergence measured via Jensen–Shannon distance between the *k*-mer spectra of partially sampled and real sequences across 128 timesteps for *k* = 7,8,9,10. Bottom: staggered score (blue) and noise schedule (black) averaged across positions and samples over timesteps (mean ± standard deviation, *n* = 1000); thin lines show staggered score individual position-averaged trajectories. Empirically defined sampling stages (exploration, consolidation, stabilization) are delimited by vertical dashed lines at *t* ≈ 31 and *t* ≈71. **b**, Top: schematic of the staggered score matrix illustrating nucleotide selection during early-stage (uniform scores, weakly directed substitutions) versus late-stage (peaked scores, directed substitutions) sampling. Bottom: normalized mutation fraction over timesteps, showing the proportion of mutations directed to the highest-scoring nucleotide (dark blue) versus equal-or lower-scoring alternatives (light blue). Total mutation counts per timestep are shown as a gray strip at the top. **c**, Paired violin plots of staggered scores at positions that undergo mutation (dark brown) versus positions that remain unchanged (light brown), shown separately for each sampling stage. Effect sizes (rank-biserial correlation *r*, Mann–Whitney U test): Stage 1 (mutation *n* = 712,056, no mutation *n* = 1,662,234) |*r*| *<* 0.05; Stage 2 (mutation *n* = 57,666, no mutation *n* = 3, 005,934) |*r*| ≈ 0.33; Stage 3 (mutation *n* = 1,495, no mutation *n* = 4,287,545) |*r*| ≈ 0.56. Effect sizes are reported to aid interpretation given the large number of observations. **d**, Single-sequence visualization of the sampling trajectory for a 230 bp lentiMPRA K562 sequence conditioned on a high activity target. Left: per-position mutation dot grid over timesteps with stage (*S*1, *S*2, *S*3) annotations. Right: Sequence logos of DeepLIFT/SHAP attribution maps (100 shuffles) color-matched to select timesteps on the mutation dot grid. Oracle-predicted activity for each partially sampled sequence is shown on the right.

This phased behavior follows the opposing interaction of the noise schedule and the staggered score as shown in Fig. 6a (bottom) and Eq. 18. The model is trained on forward-corrupted sequences to assign a score to each possible nucleotide substitution at each position. A scaled version of it (the staggered score 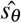) is used to sample in the reverse direction. The staggered score difference between the current (yellow boxes in Fig. 6b, top) and each candidate nucleotide measures how strongly the model favors each mutation.

Early in sampling, staggered scores are low and nearly uniform across candidate nucleotides (Fig. 6b, top) and mutations are frequent and weakly directed, encouraging broad sequence space exploration. As the noise level decreases, staggered scores become sharper and increasingly shape the transition probabilities, allowing the model to steer sequences toward activity-associated sequence features even though occasional noisy substitutions can still disrupt partially formed motif-like patterns. Late in sampling, mutations rarely happen, but among those that do, an increasing fraction land on the top-scoring nucleotide rather than equal- or lower-scoring alternatives (Fig. 6b, bottom). This is necessary to overcome the near-zero noise weighting (Eq. 18). At late timesteps, few substitutions are accepted because the noise contribution is small. Consequently, score estimates for rarely sampled alternative transitions have limited influence on the realized sequence trajectory and can appear variable without substantially affecting the final sample.

As the noise decays, mutations increasingly require stronger score support. Consistent with this behavior, the staggered score distribution at positions that mutate separates from the distribution at positions that remain unchanged by an increasing margin as sampling advances (Fig. 6c): the effect size between the two distributions (rank-biserial correlation, Mann–Whitney U test) grows monotonically, from |*r*| *<* 0.05 in Stage 1 (negligible), to |*r*| ≈ 0.33 in Stage 2 (moderate), and |*r*| ≈ 0.56 in Stage 3 (large).

Representative trajectories illustrate how these sampling dynamics coincide with the emergence of motif-associated attribution patterns (Fig. 6d; Supplementary Fig. 4). Early edits are distributed broadly across the sequence with little apparent functional structure. By the consolidation and stabilization stages, motif-like patterns emerge at high-attribution positions and oracle-predicted activity converges toward the conditioning target. The oracle serves only as a post hoc readout in this analysis: it is used to evaluate intermediate samples, but it does not enter D3’s training loss or sample selection. Thus, within an activity-conditioned sampling framework, D3 progressively constructs sequences that an independent oracle recognizes as activity-associated, without directly optimizing that oracle during sampling.

## Discussion

We developed D3 as a discrete diffusion framework for modeling and designing regulatory DNA directly in nucleotide space. Across the settings examined here, D3 provided a common discrete-diffusion framework for tasks that are often treated separately in regulatory genomics, including conditional sequence generation, experimental validation of designed regulatory sequences in K562 cells, frozen representation learning from regulatory sequence sets without activity labels, synthetic data augmentation in low-data settings, classifier-guided cell-type-conditioned generation, and constrained sequence editing through inpainting. Taken together, these results establish D3 as a flexible discrete generative framework for regulatory DNA and support its use as a platform for conditional generation, representation learning, data augmentation, and sequence design.

The comparison with genomic language models highlights the importance of the pretraining objective for regulatory sequence representation learning. In the frozen-embedding setting, D3 embeddings were more predictive than several much larger off-the-shelf genomic language model embeddings, even though D3 was trained without activity labels on a smaller, task-specific sequence set. We do not interpret this as evidence that model scale is unimportant. Rather, it suggests that the training objective matters a great deal for regulatory DNA. Genomic language models are usually trained to reconstruct nucleotide identity at every position, whereas D3 learns transition probabilities that iteratively steer sequence toward the data distribution. These results are consistent with the hypothesis that, for regulatory DNA, learning transition probabilities that move corrupted sequences toward regulatory sequence distributions can provide a more useful representation-learning signal than reconstructing nucleotide identity at every position. Distinguishing this interpretation from alternatives, including architectural inductive biases and training-data composition effects, will require direct analyses of how D3 scores vary across motif and non-motif positions in real sequences.

The experimental validation in this study is still limited, but it is encouraging. Within the training distribution, D3-generated sequences achieved measurable regulatory activity and approximated the measured activity distributions of genomic sequences in K562 lentiMPRA. This provides a proof of principle that D3-generated sequences can retain experimentally measurable regulatory activity in the assay context used for design. Even so, broader experimental validation across additional cell types, assays, and out-of-distribution design settings will be needed to define where D3 design is reliable and where it fails. Similarly, the data augmentation results should be interpreted as evidence that D3-generated sequences can improve prediction in the tested low-data setting, rather than as proof that D3 augmentation is optimal relative to all possible sequence perturbation, oversampling, or oracle-labeling strategies. Masked diffusion and masked diffusion language models are an important class of alternatives that we did not benchmark here. Comparing D3 to these formulations under matched architectures and training budgets is a natural extension of the benchmarking framework introduced here.

The current framework also has clear limitations. One is context length. D3 operates on relatively short sequence windows, which restricts its ability to model distal regulatory interactions. More efficient long-range architectures, including state-space models^42,43^, will likely be important if discrete diffusion is to scale to larger cis-regulatory contexts. A second limitation is sampling speed. D3 generates high-quality sequences by starting from random sequence and traversing the full noising schedule. That is workable for de novo generation, but inefficient for cases where the goal is to edit an existing regulatory sequence. Faster inference schemes, including formulations related to Schrodinger bridges^44^, could help shorten this trajectory. More direct editing methods that do not require full randomization would also make the framework much more useful in practice.

The mutational process is also still limited. At present it includes only substitutions. That keeps the model simple and easy to interpret, but it narrows both biological realism and design flexibility. Extending the forward and reverse processes to include insertions and deletions would better match sequence evolution and expand the range of edits the model can represent. The noise schedule is another underexplored part of the framework^45^. In D3 it has a strong effect on how broadly sampling explores sequence space and how quickly it becomes selective, so alternative schedules could affect both sample quality and efficiency. Stronger evolutionary structure in the mutation process may also improve expressivity and sampling efficiency.

D3 shows promise, but in this study it was used only as a bespoke model trained on one dataset at a time. Even under that constraint, it learned useful regulatory sequence representations and supported several design tasks. A natural next step is to ask whether the same framework can be scaled into a foundation diffusion model by training jointly across many regulatory datasets, or more broadly across genomes and species. If successful, this could yield stronger general-purpose sequence representations, broader transfer across regulatory contexts, and a more versatile generative framework for regulatory sequence design.

## Methods

### DNA Discrete Diffusion Framework

D3 adapts the Score Entropy Discrete Diffusion (SEDD) framework^30^ to operate directly in the discrete nucleotide space {*A,C, G, T*}, generating regulatory DNA sequences conditioned on specified functional activity levels or cell-type labels. Below, we describe the forward and reverse diffusion processes, the score entropy training objective, and the architectural choices underlying D3.

#### Modeling Discrete Transitions in Continuous Time

D3 employs a continuous-time Markov process over nucleotide sequences, starting from an initial distribution *q*_0_ ≈ *q*_*data*_ and driven by a transition rate matrix ℛ_*t*_ ∈ ℝ^*S*×*S*^. The process evolves according to the ODE 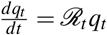, where ℛ_*t*_ has non-negative off-diagonal entries and columns summing to zero (preserving probability mass). We parameterize ℛ_*t*_ = *β* (*t*) ℛ_*b*_ with a time-dependent scalar *β* (*t*) and a fixed base rate matrix ℛ_*b*_, so that *q*_*t*_ → *p*_*re f*_ as *t* → ∞.

For small Δ*t* Euler steps for the process at any time points *t* − Δ*t* and *t*, the forward transition is:

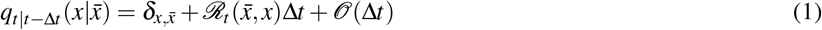

The corresponding reverse process, starting from marginal *q*_*T*_ (*x*_*T*_) and recovering *q*_*data*_(*x*_0_), uses a reverse rate matrix 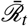:

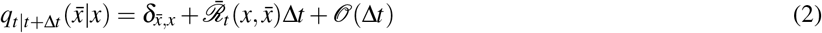

where the reverse rate satisfies:

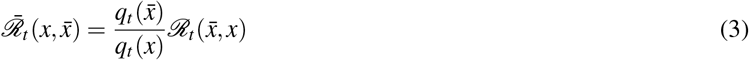

and the reverse ODE is:

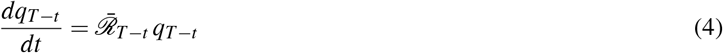

The density ratios 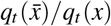 (the *concrete score*) are generally intractable but play a role analogous to ∇log *q*(*x*) in continuous diffusion, motivating learning them with a neural network.

#### Minimizing a Score Entropy Objective

Standard concrete score matching via *ℓ*_2_ loss fails because it cannot enforce positivity. SEDD instead uses score entropy, a Bregman divergence 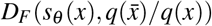 with *F*(*p*) = ∑ *p* log *p* (negative entropy), yielding the loss:

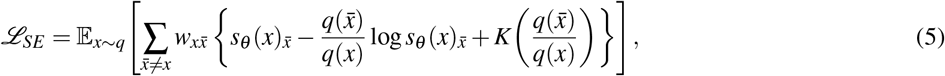

where *K*(*ℓ*) = *ℓ*(log *ℓ* − 1). The unique optimum satisfies 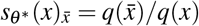 with loss value zero.

Since ℒ_*SE*_ contains the unknown ratio 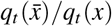, we use the denoising score entropy (DSE) formulation of^30^ (Theorem 3.4), which replaces the intractable marginal ratio with the computable perturbation kernel ratio:

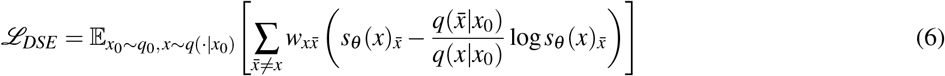

Likelihood-based training uses a time-dependent score network *s*_*θ*_ (·, *t*) with the parameterized reverse matrix 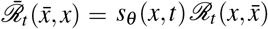. The resulting ELBO^30^ is:

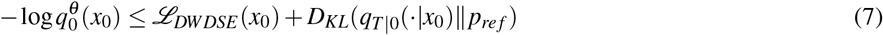

where ℒ_*DWDSE*_ (*x*_0_), the diffusion-weighted denoising score entropy, is:

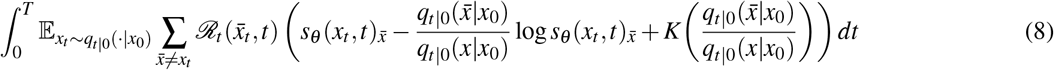

#### Choice of Forward Process

Modeling high-dimensional genomic sequences of the form **x** = *x*^1^ …*x*^*d*^ with score entropy incurs scaling challenges as *d* increases. Our state factorizes into sequences 𝒳 = {*A,C, G, T*}^*d*^ such that each position of the sequence represents one nucleotide i.e. *x*^*i*^ ∈ {*A,C, G, T*}, analogous to tokens in language models or pixels in image models. A fully general ℛ_*t*_ over sequence space would allow transitions between any pair of sequences, requiring exponential time and space, specifically 𝒪 (4^*d*^). Here, we consider a sparse structured matrix 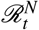 that can perturb every nucleotide independently, which is given by

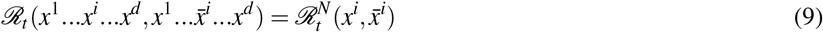

Specifically, we model all the density ratios for sequences with a Hamming distance of 1, which drastically reduces the span of perturbations to only sequences with one nucleotide change. This allows the score network to model a sequence-to-sequence map with 𝒪 (4*d*) outputs:

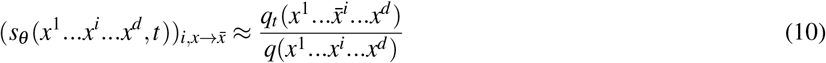

As we consider only one token change, the calculation for the forward transition 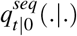 can be done in a single step by decomposing it to independent token perturbation:

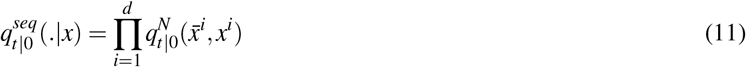

Each 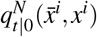 can be calculated with a noise level *β* and a pre-defined transition matrix ℛ^*N*^ as 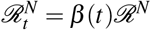. Considering 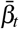 as the cumulative noise 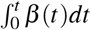, we have 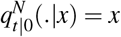 -th column of 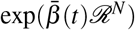. More specifically, that implies

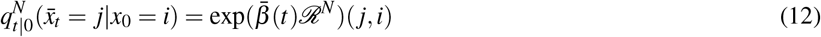

where (*j, i*) represents *j*th row, *i*th column entry of the matrix 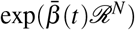. With this formulation, we follow the diffusion-weighted denoising score entropy loss ℒ_*DWDSE*_ from^30^ for our model training.

Storing all possible edge weights ℛ^*N*^(*i, j*) requires a high cost in terms of space and time. This issue can be addressed by considering special structures for transition matrix^29^. In particular, a uniform stationary base rate matrix ℛ = **11**^⊤^ − *S*ℐ can be a natural choice for categorical discrete distributions where **11**^⊤^ is a matrix of all ones, ℐ is identity matrix. For our specific scenario, the token size is 4, and hence the corresponding time-dependent transition matrix is given by

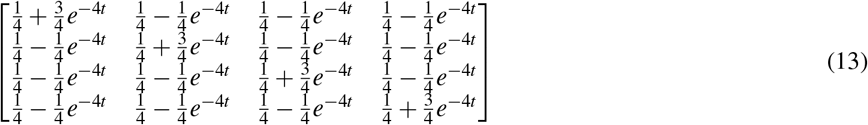

This structured matrix perturbs the input sequence according to the uniformly sampled time *t*, with the idea that all the matrix entries tend to 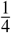 uniformly when *t* → ∞. This factorized formulation enables efficient computation of ℒ_*DWDSE*_. Specifically, ℒ_*DWDSE*_ can be calculated as below:

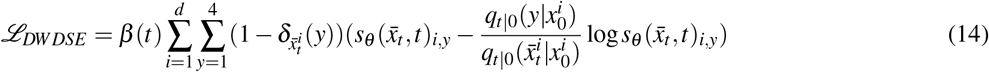

where *β* (*t*) is the noise schedule (with total noise 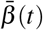), and *δ* is Kronecker delta. While other options are available, such as absorbing state process with a stationary matrix of all zeros^30^, the uniform rate matrix performed well in our experiments.

#### Choice of Generative Reverse (Sampling) Process

The parametric generative reverse process following continuous time framework with rate matrix 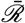 can be employed by taking a small Euler step shown in Eq. 2. While this process can be simulated using Gillespie’s Algorithm, this is impractical for large sequence lengths as stepping through each transition individually will allow only one dimension change for each backward step. Another popular approximate simulation method, tau-leaping, can be employed that applies all the transitions simultaneously. Specifically, a sequence *x*_*t*−Δ*t*_ can be constructed from *x*_*t*_ by sampling each token independently according to:

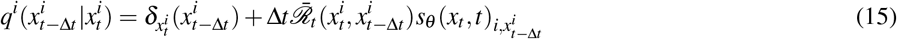

The optimal denoising can be achieved only by knowing all density ratios 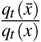. If *q*_*t*_ follows the diffusion ODE 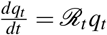, the true denoiser can be represented as a discrete version of Tweedie’s theorem, as shown in^30^

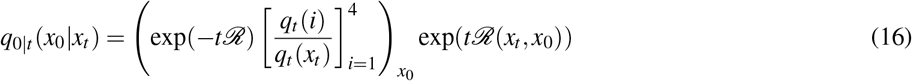

As we only know the density ratios for sequences with 1 Hamming distance away, a Tweedie denoiser analog of tau-leaping to construct transition densities can be obtained by replacing the token transition probabilities as below:

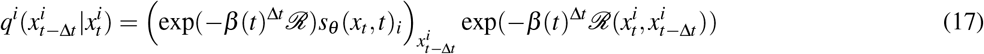

Where 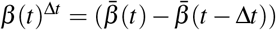. In practice, 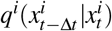 is clamped to be non-negative and normalized to sum to 1, and *x* can be obtained by sampling *i*-th token as 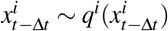.

Conditional generation is achieved by providing the conditioning signal (for example, target activity levels as scalar values) as an additional input to the score network *s*_*θ*_ (*x, t*), which enables it to learn condition-specific density ratios. However, D3 also supports sequence generation with categorical conditioning (for example, discrete cell-type labels) using classifier-free guidance (CFG), as demonstrated in the main text.

#### Mathematical Justification of the Reverse Process

The sampling process is a Bayesian inference step: given noisy state *x*_*t*_, we estimate the slightly cleaner state *x*_*t*−Δ*t*_ via

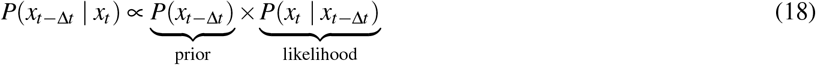

The prior corresponds to the staggered score exp(−*β* (*t*)^Δ*t*^ℛ) *s*_*θ*_ (*x*_*t*_, *t*) (the first factor in Eq. 17), where the neural network supplies discrete density-ratio estimates, 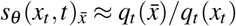, and the matrix exponential adjusts these estimates across the last diffusion step. The likelihood 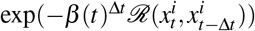 (the second factor) encodes the forward diffusion law: given *x*_*t*−Δ*t*_, how likely is the observed *x*_*t*_? Because we condition on *x*_*t*_, we evaluate this on the transposed graph, hence the name *transpose transition*. We derive both quantities below.

#### Score to Staggered Score

During sampling we step from time *t* (noise level *β*) to *t* − *dt* (noise level *β* − *dβ*). The score *s*_*θ*_ is predicted at *β*, but we need probabilities at *β* − *dβ*; the staggered score performs this adjustment via exp(− *dβ* ℛ), and 𝒮 is the number of tokens. The rate matrix for the uniform graph is

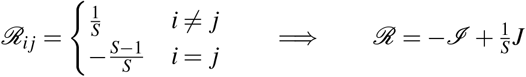

where *J* is the all-ones matrix. Its eigenstructure is: *λ*_0_ = 0 (eigenvector *v*_0_ = *S*^−1*/*2^**1**) and *λ*_1_ = −1 with multiplicity *S* − 1.

Spectral decomposition gives 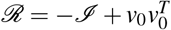, so

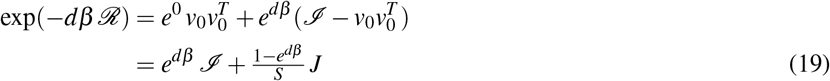

Setting *α* = *e*^−*dβ*^ this becomes

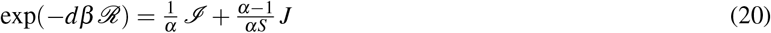

Applying this to the score vector:

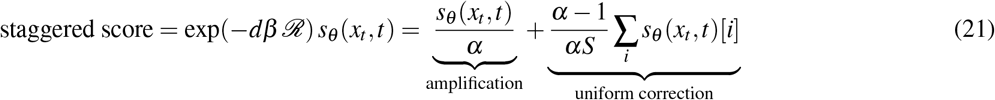

#### Transpose Transition

The forward transition matrix is 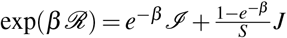 (analogous to Eq. 19), giving component-wise

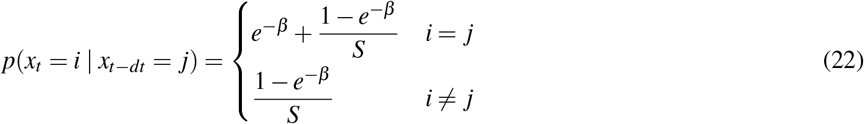

Because the uniform noise process is symmetric, the transpose transition equals the forward transition. In practice, all off-diagonal entries are initialised to 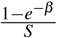 and the diagonal entry *i* is set to 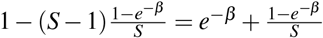.

Together, the staggered score and transpose transition provide all the components of Eq. 17 needed to generate samples with D3.

### Model Implementation

#### Diffusion Parameters

We follow the diffusion training closely from^30^ with the geometric noise distribution 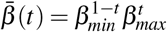 where 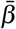 moves from *β*_*min*_ = 10^−3^ to *β*_*max*_ = 1 with increasing time step *t*. This noise schedule is chosen such that the prior loss *D*_*KL*_(*q*_*T*|0_(.|*x*_0_)||*p*_*re f*_) is negligible and 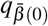 remains close to *q*_*data*_.

The uniform transition matrix is scaled down by 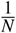 (*N* = 4 here), and we consider *p*_*re f*_ as uniform.

#### Conditioning Scheme

As mentioned in the main paper, we consider datasets with continuous-valued vectors as conditioning signals. Therefore, we send these signals with the true sequences to the score function, which is trained to learn density ratios based on the signals. For the promoter dataset, we have transcription initiation signal profiles, a vector of the same length as the sequences, i.e., 1024. Here, we concatenate the signal profiles with the sequences before forwarding to the score function. As the fly enhancer sequences have corresponding activity values in 2 contexts (housekeeping and developmental gene promoters), we employ a 2-layered fully connected network that outputs an embedding (of the same length as sequence, i.e., 249) from the 2 activity values. Then, we concatenate the embeddings with the sequences and follow the same training and sampling process.

### Architectures

#### CNN (D3-conv)

We use the same architecture for D3-Conv as the architecture used in Refs.^23,25^ for human promoter sequences. In place of the time step embedding with Gaussian random feature projection, we use similar embedding to parameterize the network with the total noise level instead of the time *t*. Here, the results (MSE) are compared with other previous methods, for which we referred to^25^. We follow a similar setup for DeepSTARR and lentiMPRA datasets. As DFM^25^ and other previous methods did not experiment with these two datasets, we trained the DFM-conv model (with a similar setup provided for the promoter) for proper performance comparison.

#### Transformer (D3-tran)

We follow the similar transformer architecture for D3-Tran as used for SEDD small^30^. We used flash attention^46^ with fused kernels and adaLN-zero time information network from^47^ with 128 hidden dimensions, which is based on incorporating time conditioning into a standard encoder-only transformer architecture^48,49^. We also use rotary positional embeddings^50^ for performance improvement. D3-Tran has 12 transformer blocks and 12 attention heads per block. Training D3-Tran with this transformer architecture outperformed D3-Conv on most metrics for all three datasets, i.e., promoter, DeepSTARR, and lentiMPRA.

Toward a more complete performance comparison of D3 with DFM, we trained DFM with the same transformer architecture with the only change of time step embedding (with Gaussian random feature projection) instead of embedding on noise level, as already mentioned for convolution model details. For a fair comparison, we did not change any other part of the transformer architecture. We followed a similar overall training setup to train DFM as explained in^25^. DFM-Tran performed poorly on several metrics, in some cases below dinucleotide-shuffled baselines (Supplementary Table 1). Because this comparison used matched architecture and training budget rather than a separately tuned DFM-Tran configuration, we interpret DFM-Tran as an architectural control rather than as an optimized implementation of DFM. The convolutional DFM-Conv configuration, which uses the original DFM architecture, performed competitively across benchmarks and serves as the more representative DFM baseline in this study.

#### Training Parameters

All of our models were trained with a learning rate 3 × 10 ^−4^, a batch size of 256 for both architectures. Following^30^, our gradient norm is clipped to 1 and a linear warmup schedule for the first 2500 iterations. We set the training for 300 epochs with a validation step after every 4 epochs. The best models were found after different training steps for different datasets and methods. We employ an exponential moving average of 0.9999 for all our D3-Conv and D3-Tran training. AdamW was used as the optimizer, with *β*_1_ = 0.9, *β*_2_ = 0.999, *ε* = 10^−8^, weight decay 0. Convolutional backbones (D3-Conv) train under 16-mixed precision (fp16 with GradScaler); transformer backbones (D3-Tran) train under bf16-mixed (except for D3-Tran for the promoter dataset, which was trained on fp16) .All the training was performed on a single node with 4 A100 80GB GPUs.

We do not perform hyperparameter optimization following Lou et al.^30^. The convolution architecture was chosen from the previous works^23,25^ and the transformer architecture from DDiT^47^, with the rotary embedding as was used in^51^. The learning rate of 3 × 10^−4^ and the EMA of 0.9999 were selected as a standard choice from the previous well-known diffusion training recipes.

### Evaluation Framework

We evaluate generated regulatory DNA sequences along three complementary dimensions: (1) *functional similarity*, (2) *compositional similarity*, and (3) *synthetic-sequence realism and novelty*.

Let *x*_real_ denote a real sequence from the dataset and *x*_gen_ a generated sequence. An oracle model *f*_*θ*_ (a pretrained deep neural network) provides an *in silico* prediction *ŷ* = *f*_*θ*_ (*x*) estimating functional activity for any input sequence *x*. We denote the conditional target activity as *y*_cond_, and true experimental measurements (if available) as *y*_exp_. Since experimental assays are costly, we rely on the oracle to approximate *y*_exp_.

Functional similarity evaluates whether generated sequences achieve functional activities similar to real sequences or the conditioning target *y*_cond_. Sequence similarity quantifies novelty and memorization relative to training data. Compositional similarity assesses whether generated sequences recapitulate k-mer spectra observed in real genomic sequences. Synthetic-sequence realism and novelty are assessed by discriminability from genomic sequences and by exact-match similarity to the training set.

#### Conditional generation fidelity

This metric measures how well generated sequences achieve the conditioning target. Given oracle predictions *ŷ*_*i*_ = *f*_*θ*_ (*x*_gen,*i*_), fidelity is the mean squared deviation from *y*_cond_ over *N* generated sequences, Fidelity 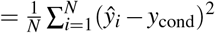; lower values indicate higher fidelity.

#### Label shift

This metric uses the Kolmogorov–Smirnov statistic *D*_KS_ = sup_*y*_ |*F*_gen_(*y*) − *F*_real_(*y*) | to compare the ECDFs of oracle predictions for *x*_gen_ and *x*_real_; lower *D*_KS_ implies more similar predictive distributions.

#### Covariate shift

This metric compares *k*-mer frequency distributions between *x*_gen_ and *x*_real_ via Jensen–Shannon divergence, 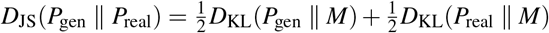, where 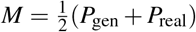. We compute 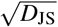 (the JS distance) for *k* = 6.

#### Discriminability

This metric quantifies the ability to distinguish generated sequences *x*_gen_ ∼ *p*_gen_ from real sequences *x*_real_ ∼ *p*_real_ by training a binary classifier *h* : *x* ↦ {0, 1} to predict whether a sequence is generated (*y* = 1) or real (*y* = 0). Discriminability is measured by AUROC, where a value near 0.5 indicates indistinguishability and a value near 1 indicates strong separability.

#### Informativeness

This metric measures whether generated sequences improve predictive power when used for dataset augmentation. Let 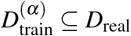 be a real training subset of fraction *α*, and let *D*_gen_ be the generated set. A supervised model *f*_*θ*_ is trained on the hybrid dataset 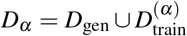 and evaluated on a held-out validation set *D*_val_. The improvement in validation metric *M* (e.g., *R*^2^, AUROC, MSE) is:

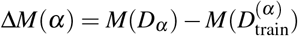

Positive Δ*M*(*α*) indicates that generated sequences contribute additional predictive information beyond the real training data alone.

#### Memorization

We identified the longest exact contiguous match between each query sequence and a reference sequence set using `BLASTN`. A nucleotide database was first constructed from the reference sequences using `makeblastdb`. Query sequences were aligned against this database with `blastn` configured to report **100% identity ungapped alignments** (`-perc_identity 100`, `-ungapped`, `-dust no`, `-soft_masking false`, `word_size=7`). All hits were retained and sorted by alignment length, and for each query the **longest match** was selected. Queries with no detected alignment were assigned an overlap length of zero so that all sequences were represented in the final table.

#### Fréchet Biological Distance (FBD)

We use FBD to measure the distributional similarity between generated and real sequences in the embedding space of a pretrained biological sequence classifier^25^. Given real sequences *x*_real_ and generated sequences *x*_gen_, each set is embedded by extracting penultimate-layer activations from the classifier, yielding embedding distributions with means *µ*_real_, *µ*_gen_ and covariances Σ_real_, Σ_gen_. FBD is then computed as:

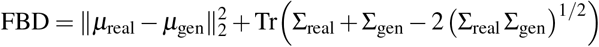

Lower FBD indicates closer alignment between generated and real sequence distributions in biologically meaningful feature space. Unlike sequence-level metrics, FBD captures higher-order functional similarity by operating in a learned representation that reflects regulatory sequence properties. We use the pretrained enhancer classifier from^25^ trained on the fly brain dataset (81 cell-type labels) and compute FBD between 10,400 held-out test sequences and an equal number of generated sequences matched by cell-type condition.

### Experimental MPRA validation in K562

#### Generation of MPRA library

The MPRA library (total of 2760 elements) design consisted of 1500 regulatory elements from the K562 large-scale library of Agarwal et al. (2025)^33^, 420 elements derived from the DFM model (synth_dfm), 420 elements derived from D3 transformer model (synth_D3 tran), and 420 elements derived from the D3 convolution model (synth_D3_conv). This MPRA library was constructed using primers and conditions as previously described^5,33^.

Briefly, the oligonucleotide pool (synthesized by Twist Bioscience, Inc.) was amplified with primers pLSmP-enh-f and minP-enh-r for 5 cycles, and the amplicons were purified with AMPure beads. Purified amplicons were amplified for an additional 15 cycles using primers (pLSmP-enh-f and pLSmP-bc-primer-r) to add 15bp sequences of random barcodes. Purified secondary amplicons were inserted into SbfI/AgeI restriction sites of the pLS-SceI vector (Addgene, Cat#137725) using NEBuilder HiFi DNA Assembly mix (NEB Cat# E2621L), followed by transformation into NEB 10-beta electrocompetent cells (NEB Cat# C3020K). Approximately 4.8 million colonies were collected from LB/carbenicillin agar plates, and plasmid DNA was purified from scraped bacteria cells using Qiagen Endo-free Plasmid Maxi columns (Qiagen, Cat# 12362). This size library suggests an average of approximately 1500 barcodes per regulatory element. The library was subjected to Next-generation sequencing (NGS) using primers^33^ containing the Illumina flow cell adapter (P5-pLSmP-ass-i17 and P7-pLSmP-ass-gfp). Library preparation procedures were based on Gordon et al. (2020)^5^. The purified amplicons from the NGS samples were sequenced using the Nextseq 2000 sequencer with the P1-XLEAP 300 cycles kit. Custom sequencing primers from Agarwal et al., 2025^33^ (pLSmP-ass-seq-R1, Read1; pLSmP-ass-seq-ind1, Barcode Read; pLSmP-ass-seq-R2, Read2) were used to perform a sequencing run consisting of paired-end 150 cycles for the regulatory sequence and an Index read of 15 cycles for the barcode.

#### Cell culture, lentivirus packaging and titration, and infections into K562 cells

The K562 cell line (ATCC, CCL-243) used for the MPRA screens was part of the Cold Spring Harbor Laboratory (CSHL) Cell Line Repository, in which all cell lines in the collection are authenticated (by Short Tandem Repeat assay) and have non-detectable levels of mycoplasma contamination. Lentivirus packaging was performed using the 293FT cell line (Thermo Fisher Scientific, Cat# R70007), which is also part of the CSHL Cell Line Repository. The culture media used to propagate the K562 cells consist of RPMI 1640 medium (Thermo Fisher Scientific, Cat# 61870127), 10% Fetal Bovine Serum (Corning, Cat# 35-010-CV), and 1X penicillin-streptomycin (Thermo Fisher Scientific, Cat# 15140122). The culture media used to grow the 293FT cell line contains DMEM (Thermo Fisher Scientific, Cat# 11995073), 10% Fetal Bovine Serum, and 1X penicillin-streptomycin Lentivirus production was carried out according to procedures described in Gordon et al., 2020^5^ and Agarwal et al., 2025^33^. After lentivirus production, the collected viral media was concentrated 30X using the Lenti-X concentrator reagent (Takara Bio, Cat# 631232) according to the manufacturer’s procedures. Concentrated virus for the MPRA validation library was frozen in aliquots and stored at -80F. To determine the amount of library lentivirus needed to achieve optimal multiplicity of infection (MOI) for high-copy screening in K562 cells, 100,000 cells per well in 24-well plates were seeded and incubated for 1 day. Serial volumes of concentrated virus (0 ul, 2 ul, 4 ul, 8 ul, 16 ul, 32 ul, 64 ul) were used to infect each well, and three replicates were set up for each infection. After 48 hours of infection, the viral media was removed from each well and replaced with fresh K562 growth media (without lentivirus). By the end of the third day post-infection, genomic DNA was extracted from all wells using the Wizard SV genomic DNA purification kit (Promega, Cat# A2361). Quantitative PCR was set up using primers that amplify viral DNA (WPRE region, forward; 5’-TACGCTGCTTTAATGCCTTTG-3’, reverse; 5’-GGGCCACAACTCCTCATAAAG-3’), plasmid backbone DNA (BB region, forward; 5’-TGCCGCATAGTTAAGCCAGTA-3’, reverse; 5’-TCAAGCCTTGCCTTGTTGTAG-3’), and the intronic region of the LIPC gene as normalization control (LIPC region, forward; 5’-TCCTCCGGAGTTATTCTTGGCA-3’, reverse; 5’-CCCCCCATCTGATCTGTTTCAC-3’). All experimental procedures and reagents for virus production and titration, and qPCR are outlined in Gordon et al. 2020^5^.

#### Lentiviral infection for MPRA screens and DNA and RNA barcode sequencing

K562 cells were propagated in T75 and T175 flasks, and 15 cm cell culture plates. Three biological replicates of screening were undertaken. For each replicate, 20 million K562 cells (infected at an estimated MOI of 10 measured by qPCR of genomic DNA after serial infections of concentrated library viruses outlined in the previous section) were seeded into a 15cm plate and incubated overnight, the next day (16 hours post-seeding cells) 1.7ml of concentrated virus and 25 ul of Polybrene (final concentration of 8 ug/ml) were added to the cells and media (with gentle and thorough mixing all components of final volume of 25 ml) for infection. After 48 hours of infection, cells were spun down, resuspended in 25ml of fresh media, and incubated at 37°C/5 % CO2 for 3 days. At the end of the third day post-media change period, genomic DNAs and RNAs were extracted from cell samples for each replicate using the Qiagen Allprep DNA/RNA Mini kit (Qiagen Inc., Cat# 80204) as per the manufacturer’s procedures. Genomic DNA and RNA samples were quantified using a Qubit fluorometer and a Bioanalyzer, respectively. NGS libraries for DNA and RNA samples were created using PCR primers from Agarwal et al., 2025^33^ and methods from Gordon et al., 2020^5^. NGS samples were sequenced using the Nextseq 2000 sequencer with the P2-XLEAP 100 cycles kit. Custom sequencing primers from Agarwal et al., 2025 (pLSmP-5bc-seq-R1, Read1; pLSmP-5bc-seq-R2, Sample Index Read; pLSmP-UMI-seq, Read2; pLSmP-bc-seq, Read3) were used to undertake a 26 cycles / 10 cycles / 10 cycles / 15 cycles run using the P2-XLEAP flowcell.

#### MPRAflow data processing pipeline

The MPRAflow 1.0 pipeline tool was installed onto the CSHL HPC cluster as per instructions according to the Shendure Lab github site (https://github.com/shendurelab/MPRAflow.git) and Gordon et al, 2020^5^. Barcodes association to MPRA validation library cis-regulatory elements. Fastq files were generated for each individual custom sequencing primer: Read1_Enhancer-forward_fastq; Read2_Enhancer-reverse_fastq; Index_Barcode_fastq. These files were used to associate barcodes with elements using the association utility in MPRAflow_1.0^5^. The final output file was a filtered_coords_to_barcodes.pickle file mapping regulatory elements to barcodes. The mean number of barcodes (filtered barcodes) associated with a given element in the library was 2017. DNA and RNA replicates barcode counting and normalization for regulatory activity measurements (RNA/DNA activity scores). Fastq files were generated for each individual custom sequencing primer: Read1_Barcode-forward_fastq; Index_Sample_fastq; Read2_UMI_fastq; Read3_Barcode-reverse_fastq. The MPRAflow_1.0 count utility was used to process the fastq files for all DNA and RNA MPRA screen samples. Briefly, for each replicate, only the subset of barcodes observed in both DNA and RNA samples for the replicate was considered. Activity scores [log2(RNA count/DNA count)] were determined and normalized for each element. A total of 91.8% (2534/2760) of regulatory elements from the library across all 3 replicates were retained for downstream analysis, with per-element barcode counts ranging from 4 to 5110. The retained set comprises 1,326 genomic K562 regulatory elements from Agarwal et al., 400 D3-transformer, 402 D3-convolutional, and 406 DFM (conv). Pairwise replicate-to-replicate Pearson correlations of log_2_(RNA*/*DNA) activity scores were *r* ≥ 0.998 across all three replicate pairs (Supplementary Fig. 5), confirming high reproducibility of the assay.

### Datasets and Oracles

#### Human Promoters

The human promoter dataset is an established benchmark for DNA generative models^23^. It contains 100, 000 promoter sequences (1024 bp, centered at the transcription start site — TSS) with their transcription initiation signal profile (measured via CAGE-seq^52^ from the FANTOM database^53^). These profiles (**p**) indicate the likelihood of transcription initiation at each base pair (**p** ∈ ℝ^1024^), which vary across genes and depend on underlying promoter sequence features. The modeling task is to generate promoter sequences conditioned on a target transcription initiation signal profile. For training, we use all 100, 000 sequences split into validation (chromosome 10; 3933 seqs), test (chromosomes 8,9; 7497 seqs) and train (all other chromosomes; 88570 seqs) sets. For sampling and evaluation, we condition on the same test set (chromosomes 8,9; 7497 seqs). Note that we do not further rank the TSSs and subset the top 40, 000 sequences as done by Aydeyev et al.^23^.

The oracle was Sei^37^, a multi-task CNN that predicts 21,907 chromatin profiles across *>* 1, 300 cell lines/tissues from DNA sequences. Active promoter predictions were obtained from the H3K4me3 chromatin mark prediction and averaged over the sequence length to obtain a per-sequence activity scalar, following Ref.^23^. The Sei oracle and weights were acquired from the DDSM repository (https://github.com/jzhoulab/ddsm).

#### Context-specific Fly Enhancers

The fly enhancer dataset contains *D. melanogaster* S2 cell enhancer sequences with corresponding activity values across two transcriptional programs: housekeeping and developmental. We retrieve it from the STARR-seq dataset by Almeida et al.^32^. Each enhancer sequence (249 bp) has two scalar activity values for the HK, DEV conditions split across train (402, 296 seqs), validation (40, 570 seqs) and test (41, 186 seqs) sets. Sequences containing N characters were removed, which had a minimal impact on the dataset size (reducing it by approximately 0.005%). The oracle was a DeepSTARR model trained with EvoAug^54^, as it demonstrated improved performance over the original training.

#### Human CRE activity (lentiMPRA)

We compile a dataset of human cis-regulatory elements (CREs) with previously reported cell-type specific activity measured via lentivirus-based massively parallel reporter assays (lentiMPRAs)^33^. In short, these assays inserted thousands of (candidate) human CREs (enhancers and promoters) into reporter plasmid libraries with barcodes, which were then integrated into two cell lines (K562: lymphoblasts, HepG2: hepatocytes). CRE activity was then quantified by sequencing transcribed RNA barcodes. We use the large-scale single-cell-type libraries for our benchmark (226, 255 sequences for K562 and 139, 886 sequences for HepG2), where we use the larger version of the K562 library as in^27^.

The oracle was an MPRA-LegNet model^55^, an EfficientNet-inspired convolutional architecture with squeeze-and-excitation (SE) attention. The model takes one-hot encoded DNA sequences (4 × 230) and outputs a scalar. The model was trained for 25 epochs using AdamW (weight decay 0.1) with a OneCycleLR schedule (max learning rate 0.01, 30% warmup), gradient clipping at 1.0, mixed precision (FP16), and a batch size of 1024. Data augmentation included reverse complement and sequence shift augmentation. A separate model was trained for each cell line.

## Acknowledgments

We thank members of the Koo Lab for helpful comments on the manuscript. We are grateful to Surag Nair and Gökçen Eraslan for helpful discussions and feedback on this work. This research was supported in part by the National Human Genome Research Institute of the National Institutes of Health under Award Number R01HG012131 and the National Institute of General Medical Sciences of the National Institutes of Health under Award Number R01GM149921. This work was performed with assistance from the US National Institutes of Health Grant S10OD028632-01. The Simons Center for Quantitative Biology at Cold Spring Harbor Laboratory is supported by a generous endowment from the Simons Foundation. The content is solely the responsibility of the authors and does not necessarily represent the official views of the National Institutes of Health.

## Author contributions

A.S. and P.K. conceived the study and designed the overall framework. A.S. developed the D3 model, implemented the training and sampling code, and led computational experiments. A.S. contributed to the classifier-free guidance experiments and led the representation learning and linear probing analyses; J.Z. and P.M. contributed to generating the associated benchmark results for linear probing analyses. Y.Y. restructured the codebase and generated visualizations that aided mechanistic understanding of the generative process. Z.T. and P.K. contributed to evaluation framework design and benchmarking. N.S. and Y.K. contributed to the data augmentation and low-data regime experiments; A.D. rigorously performed all experiments to finalize the results. A.D. and P.K. designed the inpainting experiments, which A.D. executed. Y.Y. and M.N. contributed to data processing and computational benchmarking for the wet-lab lentiMPRA validation experiments; K.H. and K.C. designed and performed the wet-lab validation experiments. D.L., A.D., and A.S. contributed to the mechanistic understanding and interpretation of the generative process. A.D., A.S., D.L., and Y.Y. planned the figures with input from P.K., and A.D. prepared them. A.S., A.D., and P.K. wrote the manuscript with input from all authors. P.K. supervised the project, secured funding, and provided overall direction.

## Competing interests

P.K. is a paid consultant for Inari Agriculture, Inc. and holds stock options in Inari Agriculture, Inc. P.K. serves on the Scientific Advisory Board of Collage Bio Inc. and holds stock options in Collage Bio Inc. The remaining authors declare no competing interests.

## Data and materials availability

All research code is available at https://github.com/anirbansarkar-cs/D3-DNA-Discrete-Diffusion. An installable Python package for D3, supporting training, sampling, and evaluation, is available at https://github.com/anirbansarkar-cs/d3-dna. The benchmark can be reproduced with the metrics in the package. Trained model weights, oracle models, and datasets used in this study are deposited at Zenodo (https://zenodo.org/records/19774653).

---

**Table S1.** Tabulated D3 benchmark across three regulatory genomics datasets. Reported values correspond to the bar plots in Figure 2. Best result per column is in bold.

| Metric | Method | CAGE | STARR-seq | lentiMPRA-K562 | lentiMPRA-HepG2 |
| --- | --- | --- | --- | --- | --- |
| MSE ↓ | Shuffle | 0.040 | 1.688 | 0.373 | 0.722 |
|  | DFM (tran) | 0.034 | 2.231 | 0.311 | 0.580 |
|  | DFM (conv) | 0.029 | 1.228 | 0.245 | 0.476 |
|  | DDSM | 0.028 | — | — | — |
|  | D3 (conv) | **0.027** | 1.133 | 0.208 | 0.466 |
|  | D3 (tran) | **0.027** | **1.083** | **0.200** | **0.446** |
| KS Statistic ↓ | Shuffle | 0.301 | 0.206 | 0.431 | 0.316 |
|  | DFM (tran) | 0.176 | 0.277 | 0.231 | 0.717 |
|  | DFM (conv) | 0.095 | 0.033 | 0.164 | 0.086 |
|  | DDSM | 0.049 | — | — | — |
|  | D3 (conv) | 0.067 | 0.029 | 0.056 | 0.045 |
|  | D3 (tran) | **0.045** | **0.022** | **0.008** | **0.043** |
| JS Distance ↓ | Shuffle | 0.096 | 0.085 | 0.185 | 0.189 |
|  | DFM (tran) | 0.135 | 0.211 | 0.195 | 0.831 |
|  | DFM (conv) | 0.087 | 0.047 | 0.054 | 0.051 |
|  | DDSM | 0.042 | — | — | — |
|  | D3 (conv) | **0.024** | **0.031** | 0.034 | 0.054 |
|  | D3 (tran) | 0.029 | 0.034 | **0.023** | **0.022** |
| AUROC ↓ | Shuffle | 0.994 | 0.945 | 1.000 | 1.000 |
|  | DFM (tran) | 0.988 | 0.992 | 0.994 | 1.000 |
|  | DFM (conv) | 0.870 | 0.678 | 0.757 | 0.790 |
|  | DDSM | 0.683 | — | — | — |
|  | D3 (conv) | 0.560 | **0.610** | 0.775 | 0.791 |
|  | D3 (tran) | **0.553** | 0.647 | **0.727** | **0.765** |
| Match Fraction | Shuffle | 0.019 | 0.069 | 0.072 | 0.072 |
|  | Genomic (test) | 0.025 | 0.077 | 0.111 | 0.107 |
|  | DFM (conv) | 0.021 | 0.070 | 0.107 | 0.105 |
|  | D3 (tran) | 0.025 | 0.076 | 0.102 | 0.106 |

**Figure S1.**
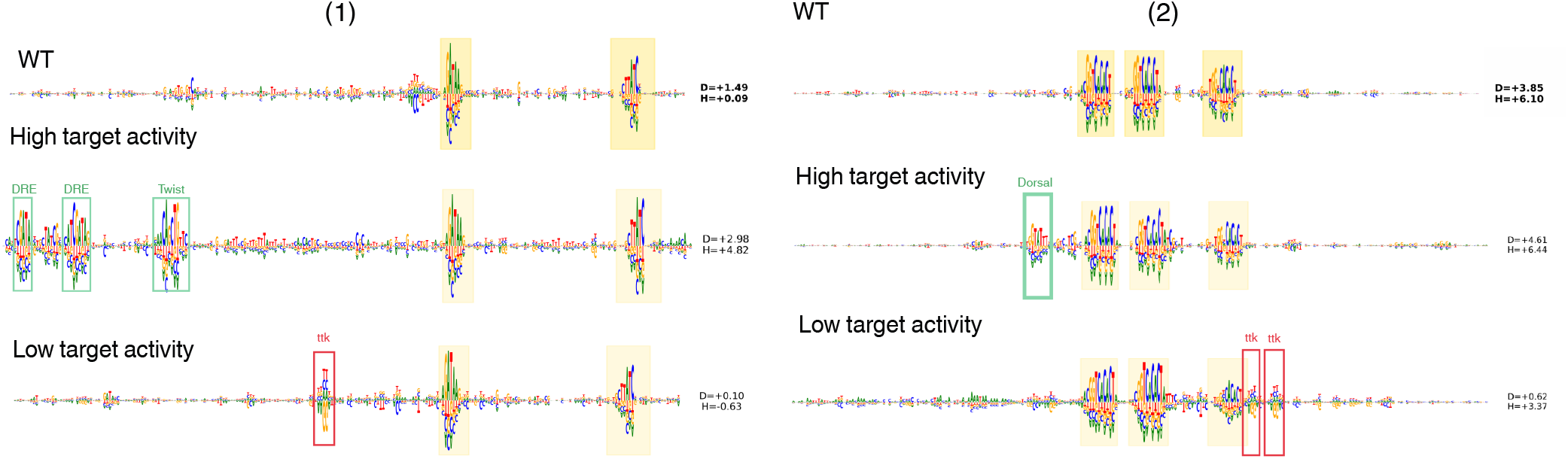
D3 inpainting can generate sequence context around fixed motif instances. Attribution maps (DeepLIFT/SHAP) from the DeepSTARR developmental head for two additional high-activity wild-type sequences conditioned on high or low activity targets (yellow shading marks fixed motif regions) show that high-activity conditioning recovers known activating motifs — DRE, Twist, and Dorsal — while low-activity conditioning surfaces ttk repressor signatures and AAAGA 6-mers.

**Figure S2.**
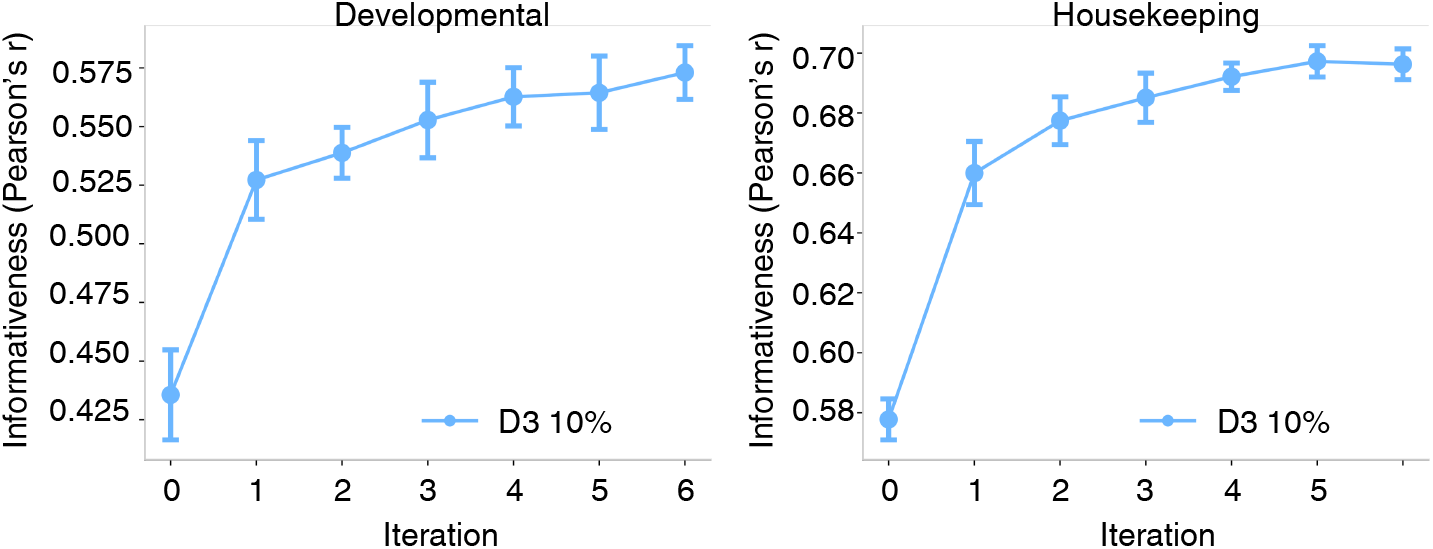
Plot of performance for DeepSTARR models trained on the 10% subset augmented with D3-generated sequences at increasing augmentation sizes (1× –6× the real subset size) for developmental and housekeeping tasks. Pearson’s *r* between predicted and measured enhancer activity on the STARR-seq test set. Error bars represent one standard deviation across three independent training runs.

**Figure S3.**
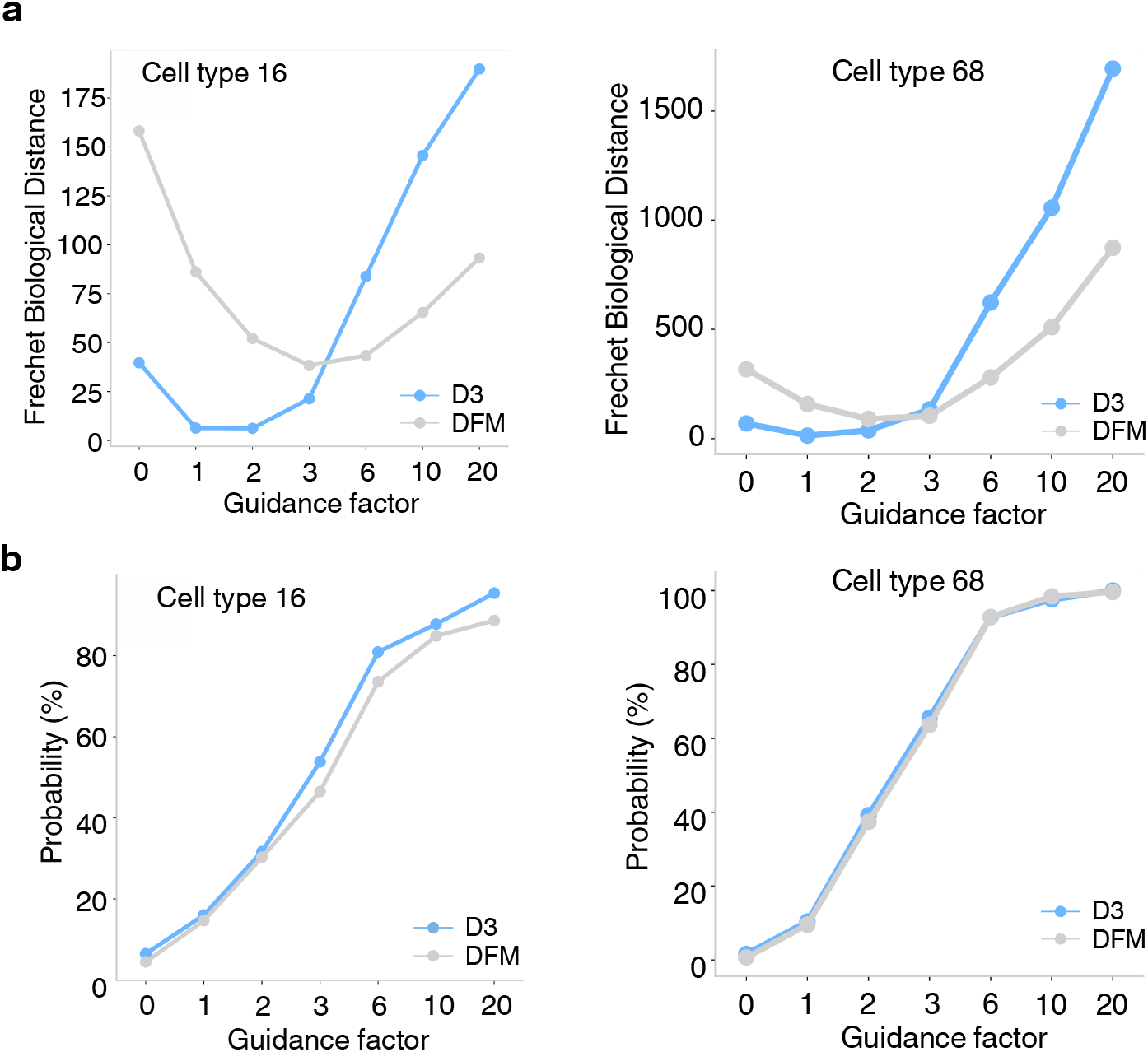
Additional examples of classifier-free guidance. **a**, Fréchet Biological Distance (FBD) between generated and held-out real sequences as a function of CFG guidance strength, for D3 and DFM. Sequences are generated conditioned on cell types 16 and 68 labels from the *Drosophila* brain enhancer dataset. FBD is computed in the embedding space of a pretrained *Drosophila* brain enhancer classifier; lower values indicate closer alignment to the real sequence distribution. **b**, Target cell-type classification probability assigned by a pretrained enhancer classifier to sequences generated for cell type 16 and 68 as a function of guidance strength for D3 and DFM.

**Figure S4.**
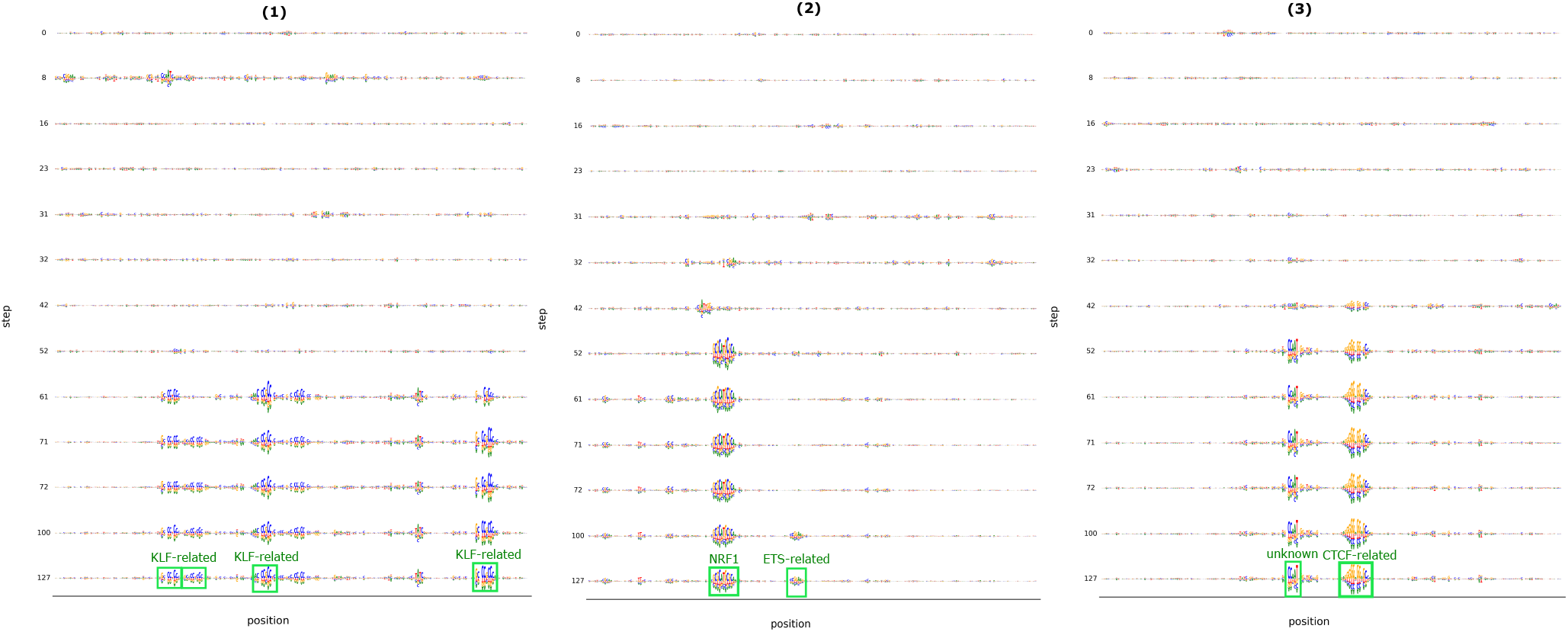
Three representative examples of DeepLIFT/SHAP attribution maps of intermediate lentiMPRA K562 sampled sequences at select timesteps, for high activity targets. Sequences evolve from a random-like state (top) to a functional state (bottom). D3-sampled sequences recover motifs (green boxes) that qualitatively resemble TF-MoDISco motifs identified by Agarwal et al. using MPRALegnet as an oracle directly on genomic sequences of K562 cells: KLF-related, CTCF-related, and NRF1 motifs, as well as similar unknown attribution patterns.

**Figure S5.**
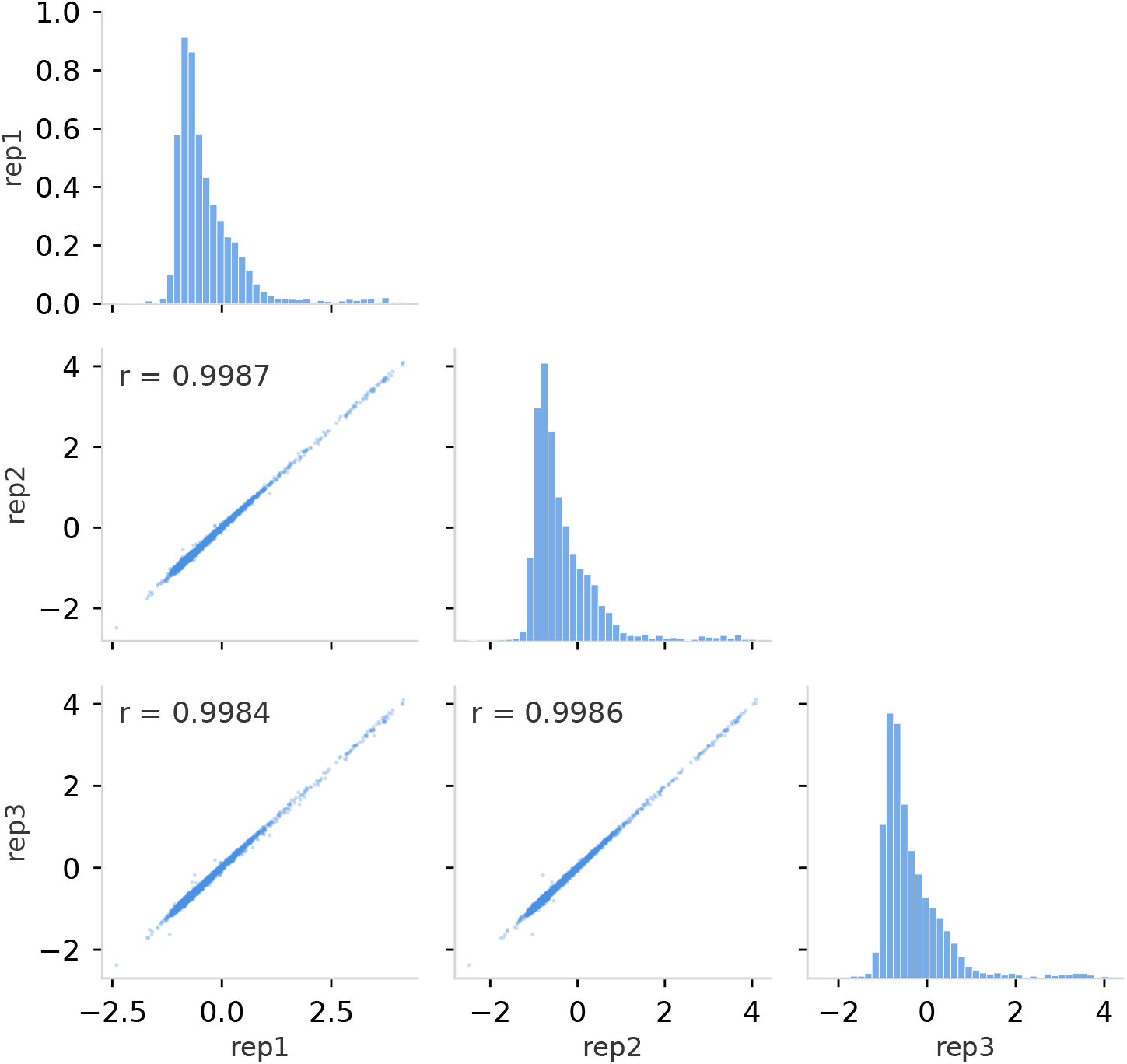
Correlation analysis of technical replicates for lentiMPRA experiments. Pairwise concordance of lentiMPRA technical replicates for enchancer activity defined as log_2_(RNA*/*DNA) for the *N* = 2,774 cis-regulatory sequence elements (87.23% of the 3,180-element library) that passed barcode-count filtering in all three biological replicates. Retained elements comprise 1,326 genomic K562 sequences from Agarwal et al. (2025), 1,448 synthetic sequences. Diagonal panels show the marginal distribution of log_2_(RNA*/*DNA) per replicate. Lower-triangle panels show scatter plots across different pairs of replicates, where each dot represents a different sequence. Pearson correlation coefficient annotated in the upper-left corner (rep1 vs rep2: Pearson’s *r* = 0.999; rep1 vs rep3: Pearson’s *r* = 0.998; rep2 vs rep3: Pearson’s *r* = 0.999).

## Notes

### Summary of Updates

Updated the experiments, analysis, and presentation.

## References

1. Kim, S. & Wysocka, J. Deciphering the multi-scale, quantitative cis-regulatory code. Mol. cell 83, 373–392 (2023).

2. Tewhey, R. et al. Direct identification of hundreds of expression-modulating variants using a multiplexed reporter assay. Cell 165, 1519–1529 (2016).

3. Sahu, B. et al. Sequence determinants of human gene regulatory elements. Nat. Genet. 54, 283–294 (2022).

4. Arnold, C. D. et al. Genome-wide quantitative enhancer activity maps identified by starr-seq. Science 339, 1074–1077 (2013).

5. Gordon, M. G. et al. lentimpra and mpraflow for high-throughput functional characterization of gene regulatory elements. Nat. protocols 15, 2387–2412 (2020).

6. Koo, P. K. & Ploenzke, M. Deep learning for inferring transcription factor binding sites. Curr. opinion systems biology 19, 16–23 (2020).

7. Barbadilla-Martínez, L., Klaassen, N., van Steensel, B. et al. Predicting gene expression from dna sequence using deep learning models. Nat. Rev. Genet. DOI: 10.1038/s41576-025-00841-2 (2025).

8. Linder, J. & Seelig, G. Fast differentiable dna and protein sequence optimization for molecular design. arXiv preprint 2005.11275 (2020).

9. Schreiber, J., Lu, Y. Y. & Noble, W. S. Ledidi: Designing genomic edits that induce functional activity. bioRxiv 2020.05.21.109686, DOI: 10.1101/2020.05.21.109686 (2020).

10. Gosai, S. J. et al. Machine-guided design of synthetic cell type-specific cis-regulatory elements. bioRxiv (2023).

11. Vaishnav, E. D. et al. The evolution, evolvability and engineering of gene regulatory dna. Nature 603, 455–463 (2022).

12. Taskiran, I. I. et al. Cell-type-directed design of synthetic enhancers. Nature 626, 212–220 (2024).

13. Jain, M. et al. Biological sequence design with gflownets. In International Conference on Machine Learning, 9786–9801 (PMLR, 2022).

14. de Almeida, B. P. et al. Targeted design of synthetic enhancers for selected tissues in the drosophila embryo. Nature 626, 207–211 (2024).

15. Castillo-Hair, S. M. et al. Programming human cell type-specific gene expression via an atlas of AI-designed enhancers. bioRxiv 2025.09.30.679565, DOI: 10.1101/2025.09.30.679565 (2025).

16. Yin, C. H. et al. Iterative deep learning design of human enhancers exploits condensed sequence grammar to achieve cell type-specificity. Cell Syst. 16, 101302, DOI: 10.1016/j.cels.2025.101302 (2025).

17. Krützfeldt, L.-M., Schubach, M. & Kircher, M. The impact of different negative training data on regulatory sequence predictions. PLOS ONE 15, e0237412, DOI: 10.1371/journal.pone.0237412 (2020).

18. Rohs, R. et al. The role of dna shape in protein–dna recognition. Nature 461, 1248–1253 (2009).

19. Mariani, L. et al. DNA bendability regulates transcription factor binding to nucleosomes. Nat. Struct. & Mol. Biol. 32, 2185–2195, DOI: 10.1038/s41594-025-01633-2 (2025).

20. Zhou, Z. et al. Dnabert-2: Efficient foundation model and benchmark for multi-species genome. arXiv (2023).

21. Dalla-Torre, H. et al. Nucleotide transformer: building and evaluating robust foundation models for human genomics. Nat. Methods 22, 287–297 (2025).

22. Brixi, G. et al. Genome modeling and design across all domains of life with evo 2. BioRxiv 2025–02 (2025).

23. Avdeyev, P., Shi, C., Tan, Y., Dudnyk, K. & Zhou, J. Dirichlet diffusion score model for biological sequence generation. In International Conference on Machine Learning, 1276–1301 (PMLR, 2023).

24. DaSilva, L. F. et al. Designing synthetic regulatory elements using the generative ai framework dna-diffusion. Nat. Genet. 1–15 (2025).

25. Stark, H. et al. Dirichlet flow matching with applications to dna sequence design. arXiv preprint 2402.05841 (2024).

26. Benegas, G., Ye, C., Albors, C., Li, J. C. & Song, Y. S. Genomic language models: opportunities and challenges. Trends Genet. (2025).

27. Tang, Z. & Koo, P. K. Evaluating the representational power of pre-trained dna language models for regulatory genomics. bioRxiv 2024–02 (2024).

28. Patel, A. et al. Dart-eval: A comprehensive dna language model evaluation benchmark on regulatory dna. Adv. Neural Inf. Process. Syst. 37, 62024–62061 (2025).

29. Austin, J., Johnson, D. D., Ho, J., Tarlow, D. & Van Den Berg, R. Structured denoising diffusion models in discrete state-spaces. Adv. Neural Inf. Process. Syst. 34, 17981–17993 (2021).

30. Lou, A., Meng, C. & Ermon, S. Discrete diffusion language modeling by estimating the ratios of the data distribution. arXiv preprint 2310.16834 (2023).

31. Jukes, T. H., Cantor, C. R. et al. Evolution of protein molecules. Mammalian protein metabolism 3, 132 (1969).

32. de Almeida, B. P., Reiter, F., Pagani, M. & Stark, A. Deepstarr predicts enhancer activity from dna sequence and enables the de novo design of synthetic enhancers. Nat. genetics 54, 613–624 (2022).

33. Agarwal, V. et al. Massively parallel characterization of transcriptional regulatory elements. Nature 1–10 (2025).

34. Patel, A. et al. Dart-eval: A comprehensive dna language model evaluation benchmark on regulatory dna. arXiv preprint 2412.05430 (2024).

35. Nguyen, E. et al. Hyenadna: Long-range genomic sequence modeling at single nucleotide resolution. arXiv preprint 2306.15794 (2023).

36. Benegas, G., Batra, S. S. & Song, Y. S. Dna language models are powerful predictors of genome-wide variant effects. Proc. Natl. Acad. Sci. 120, e2311219120 (2023).

37. Chen, K. M., Wong, A. K., Troyanskaya, O. G. & Zhou, J. A sequence-based global map of regulatory activity for deciphering human genetics. Nat. genetics 54, 940–949 (2022).

38. Avsec, Ž. et al. Effective gene expression prediction from sequence by integrating long-range interactions. Nat. methods 18, 1196–1203 (2021).

39. Koo, P. K., Majdandzic, A., Ploenzke, M., Anand, P. & Paul, S. B. Global importance analysis: An interpretability method to quantify importance of genomic features in deep neural networks. PLoS computational biology 17, e1008925 (2021).

40. Toneyan, S., Tang, Z. & Koo, P. K. Evaluating deep learning for predicting epigenomic profiles. Nat. machine intelligence 4, 1088–1100 (2022).

41. Janssens, J. et al. Decoding gene regulation in the fly brain. Nature 601, 630–636 (2022).

42. Gu, A. & Dao, T. Mamba: Linear-time sequence modeling with selective state spaces. arXiv preprint 2312.00752 (2023).

43. Poli, M. et al. Hyena hierarchy: Towards larger convolutional language models. In International Conference on Machine Learning, 28043–28078 (PMLR, 2023).

44. De Bortoli, V., Thornton, J., Heng, J. & Doucet, A. Diffusion schrödinger bridge with applications to score-based generative modeling. Adv. Neural Inf. Process. Syst. 34, 17695–17709 (2021).

45. Amin, A. N., Gruver, N. & Wilson, A. G. Why masking diffusion works: Condition on the jump schedule for improved discrete diffusion. arXiv preprint 2506.08316 (2025).

46. Dao, T., Fu, D., Ermon, S., Rudra, A. & Ré, C. F. Fast and memory-efficient exact attention with io-awareness, 2022. URL https://arxiv.org/abs/2205.14135.

47. Peebles, W. & Xie, S. Scalable diffusion models with transformers. In Proceedings of the IEEE/CVF International Conference on Computer Vision, 4195–4205 (2023).

48. Vaswani, A. et al. Attention is all you need. Adv. neural information processing systems 30 (2017).

49. Devlin, J. et al. Bert: Pre-training of deep bidirectional transformers for language understanding. arXiv 1810.04805 (2018).

50. Su, J. et al. Roformer: Enhanced transformer with rotary position embedding. Neurocomputing 568, 127063 (2024).

51. Gulrajani, I. & Hashimoto, T. B. Likelihood-based diffusion language models. Adv. Neural Inf. Process. Syst. 36 (2024).

52. Shiraki, T. et al. Cap analysis gene expression for high-throughput analysis of transcriptional starting point and identification of promoter usage. Proc. Natl. Acad. Sci. 100, 15776–15781 (2003).

53. Carninci, P. et al. The transcriptional landscape of the mammalian genome. Science 309, 1559–1563, DOI: 10.1126/science.1112014 (2005).

54. Lee, N. K., Tang, Z., Toneyan, S. & Koo, P. K. Evoaug: improving generalization and interpretability of genomic deep neural networks with evolution-inspired data augmentations. Genome Biol. 24, 105 (2023).

55. Penzar, D. et al. Legnet: a best-in-class deep learning model for short dna regulatory regions. Bioinformatics 39, btad457 (2023).

